# Chemogenetic silencing of Na_V_1.8 positive sensory neurons reverses chronic neuropathic and bone cancer pain in FLEx PSAM^4^-GlyR mice

**DOI:** 10.1101/2023.08.15.553398

**Authors:** Rayan Haroun, Samuel J Gossage, Ana Paula Luiz, Manuel Arcangeletti, Shafaq Sikandar, James J Cox, Jing Zhao, John N Wood

## Abstract

Drive from peripheral neurons is essential in almost all pain states, but pharmacological silencing of these neurons to effect analgesia has proved problematic. Reversible gene therapy using long-lived chemogenetic approaches is an appealing option. We used the genetically-activated chloride channel PSAM^4^ -GlyR to examine pain pathways in mice. Using recombinant AAV9-based delivery to sensory neurons, we found a reversal of acute pain behavior and diminished neuronal activity using *in vitro* and *in vivo* GCaMP imaging upon activation of PSAM^4^ -GlyR with varenicline. A significant reduction in inflammatory heat hyperalgesia and oxaliplatin-induced cold allodynia was also observed. Importantly, there was no impairment of motor coordination, but innocuous von Frey sensation was inhibited. We generated a transgenic mouse that expresses a CAG-driven FLExed PSAM^4^ -GlyR downstream of the *Rosa26* locus that requires Cre recombinase to enable the expression of PSAM^4^ -GlyR and tdTomato. We used Na_V_1.8 Cre to examine the role of predominantly nociceptive Na_V_1.8+ neurons in cancer-induced bone pain (CIBP) and neuropathic pain caused by chronic constriction injury (CCI). Varenicline activation of PSAM^4^ -GlyR in Na_V_1.8-positive neurons reversed CCI-driven mechanical, thermal, and cold sensitivity. Additionally, varenicline treatment of mice with CIBP expressing PSAM^4^ -GlyR in Na_V_1.8+ sensory neurons reversed cancer pain as assessed by weight-bearing. Moreover, when these mice were subjected to acute pain assays, an elevation in withdrawal thresholds to noxious mechanical and thermal stimuli was detected, but innocuous mechanical sensations remained unaffected. These studies confirm the utility of PSAM^4^ -GlyR chemogenetic silencing in chronic pain states for mechanistic analysis and potential future therapeutic use.

**Significance statement:** Chronic pain is a massive problem. Peripheral nerve block is effective in many chronic pain conditions, demonstrating the importance of peripheral drive in chronic pain. We used chemogenetic tools based on the modified ligand-gated chloride channel PSAM^4^ -GlyR to silence dorsal root ganglion neurons *in vitro* and *in vivo*. This approach reduces pain-like behavior in acute and chronic pain models, including resistant pain conditions like neuropathic pain or cancer-induced bone pain. We generated a mouse line that expresses PSAM^4^ -GlyR in a Cre-dependent manner, providing a useful research tool to address not only the role of nociceptive sensory neurons in pain states but also the function of genetically defined sets of neurons throughout the nervous system in normal and pathological conditions.

## Introduction

A fifth of the global population suffers from chronic pain, and many individuals are inadequately treated (Breivik et al., 2006). As populations age, both the total number and percentage of people experiencing pain are rising (Freburger et al., 2009; Tompkins et al., 2017). The issue of pain thus constitutes potentially the biggest clinical challenge of the twenty-first century, quantitatively and qualitatively. Short-lived pain, such as post-surgical pain, is successfully managed by non-steroidal anti-inflammatory drugs (NSAIDs) and opioids, while persistent pain states are much more difficult to control (Saragiotto et al., 2016). Opioids are frequently and inappropriately prescribed for chronic pain (Stannard, 2018). Excessive opioid prescription has led to an epidemic of opioid addiction. Over 80,000 individuals died of opioid overdoses in the US alone in 2021, creating a tragedy of enormous proportions (NIDA, 2023).

Different peripheral mechanisms are involved in distinct chronic pain states. Redundancy and plasticity within the peripheral damage-sensing neurons confirm that multiple molecular targets are at play (Bangash et al., 2018). It is, therefore, attractive to exploit therapeutic approaches that target the activity of subsets of neurons rather than individual molecular targets. Reversible gene therapy involving chemogenetic approaches, such as DREADDS (Designer Receptors Activated by Designer Drugs) (Roth, 2016) and, more recently PSAMs (pharmacologically activated actuator modules) (Sternson and Bleakman, 2020), is an attractive route to silencing the sets of peripheral neurons that drive chronic pain (Magnus et al., 2011). PSAMs rely on a modified ligand binding domain of the α7 nicotinic acetylcholine receptor (nAChR) linked to the ion pore domains of anion or cation channels such as glycine or 5HT3 channels, respectively, in order to silence or activate neurons. We exploited PSAM^4^ -GlyR, which relies on a triple mutant ligand binding domain of α7-nAChR, spliced onto the ion pore domain of glycine receptors (GlyR) for neuronal silencing (Magnus et al., 2019). Activation of PSAM^4^ -GlyR is affected by treatment with the exogenous ligand varenicline, whilst normal nicotinic ligands are inactive (Magnus et al., 2019).

We used both recombinant adeno-associated virus (AAV9) to deliver PSAM^4^ -GlyR, as this has therapeutic potential, as well as a mouse line that expresses FLExed PSAM^4^ -GlyR for detailed mechanistic studies. We tested whether the silencing of nociceptive neurons that express the voltage-gated sodium channel Na_V_1.8 could reduce the pain-like behavior associated with chronic pain, including cancer-induced bone pain (CIBP) and neuropathic pain.

The involvement of Na_V_1.8+ neurons in pain is well established in animal models of mechanical and inflammatory pain, and gain-of-function mutations leading to erythromelalgia have been identified in humans (Akopian et al., 1999; Kist et al., 2016). More than 80% of nociceptive input has been attributed to action potentials linked to this channel in mice (Renganathan et al., 2001). Selective inhibition of Na_V_1.8 has been shown to inhibit both inflammatory and neuropathic pain (Ekberg et al., 2006). A role for this channel in CIBP has been established as the current density of Na_V_1.8 increases significantly in the DRG neurons following CIBP, and blockade of this channel alleviated mechanical and thermal sensitivity associated with CIBP (Miao et al., 2010; Liu et al., 2014). We, therefore, assessed the role of Na_V_1.8-positive neurons in an optimized CIBP model by means of Na_V_1.8 Cre PSAM^4^ -GlyR-based silencing. We compared the effects of neuronal silencing on bone pain behavior with mice where the Na_V_1.8 sensory neurons had been deleted using diphtheria toxin. We present evidence that PSAM^4^ -GlyR activation is a highly effective mechanism for nociceptive neuronal silencing with consequent reversal of CIBP. Chemotherapy–evoked pain using oxaliplatin was also explored. We examined neuropathic pain behavior using a chronic constriction injury model and found that PSAM^4^ -GlyR-based silencing of the Na_V_1.8+ neurons was also able to reverse neuropathic pain. The potential for both therapeutic use with viral delivery, as well as mechanistic insights into the neuronal function using Cre-dependent PSAM^4^ - GlyR mice, is discussed.

## Materials and Methods

### Cloning CMV PSAM^4^-GlyR IRES mCherry WPRE BGH pA

Plasmid A (Vectorbuilder) (see the sequence map in Figure 1) was subjected to restriction digestion using two restriction enzymes: XbaI and AccI (New England Biolabs (NEB)) in the cutsmart buffer. The restriction digestion reaction mixture was incubated at 37°C for 16 hours. Then the reaction products were run on a gel (1% agarose in 1x Tris-acetate-EDTA (TAE) buffer). The reaction gave rise to two bands: 3755 bp and 2074 bp. The band corresponding to 3755bp was extracted using the Qiagen gel extraction kit following the manufacturer’s protocol. The extracted 3755bp band will be named ‘Cut vector’ as it contained the backbone with inverted terminal repeats (ITRs). The Woodchuck hepatitis virus posttranscriptional regulatory element (WPRE) sequence and Bovine growth hormone polyadenylation signal (BGH pA) were present in this sequence.

**Figure 1:**
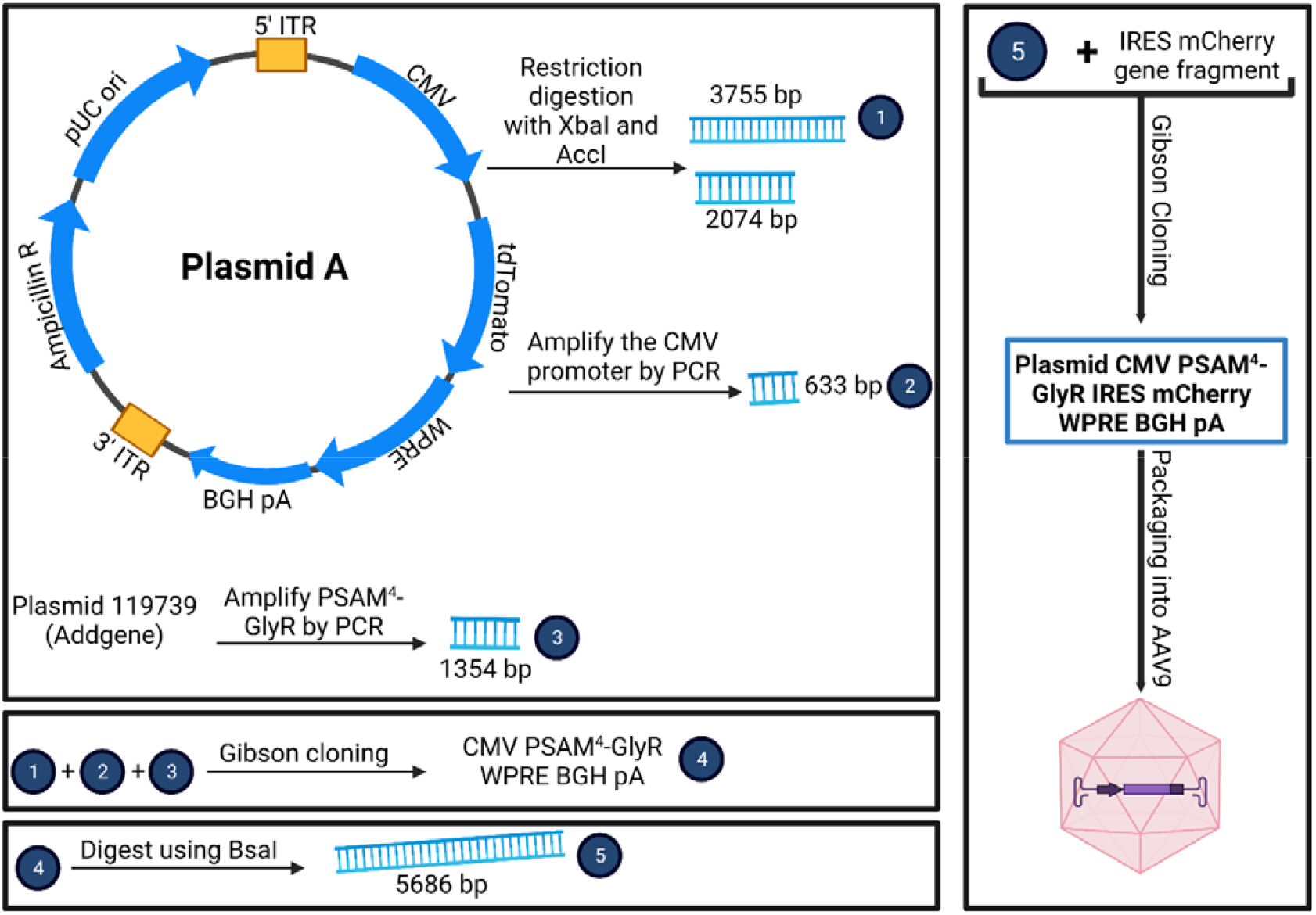
Cloning protocol for plasmid CMV PSAM^4^ -GlyR IRES mCherry WPRE BGH pA. ITR: inverted terminal repeats, WPRE: Woodchuck hepatitis virus posttranscriptional regulatory element sequence, BGH pA: Bovine growth hormone polyadenylation signal, CMV: cytomegalovirus promoter sequence, IRES: Internal Ribosome Entry Sites, PCR: polymerase chain reaction, Ampicillin R: ampicillin resistance.

To produce a CMV-driven PSAM^4^ -GlyR WPRE BGH pA sequence, the CMV promoter sequence and the PSAM^4^ -GlyR sequences needed to be amplified from other vectors to enable their cloning into the ‘cut vector’ sequence via Gibson cloning (Kalva et al., 2018). For amplifying the CMV promoter sequence, plasmid A was used as a template. The PCR protocol was 95°C 3 mins, 98°C 20 secs, 55-60°C annealing gradient for 30 secs, 72°C (1min/1kb), cycles from step 2 were repeated 34 times, 72°C for 5 mins, incubation at 12°C. The forward primer used for this PCR was (<u>TCACTAGGGGTTCCTTCTAG</u>GTATAGAAAAGTTGTAGTTATTAAT), and the reverse primer was (<u>TACAAACTTGGATCTGACGG</u>TTTCACTAGGGGTTCCTTCTAG). These two primers allowed the addition of appropriate 20-bp overhangs (underlined) on each side of the CMV promoter fragment to permit the use of the TakaraBio In-fusion kit for cloning, which necessitates the presence of at least 15 bp overhangs on each side. To amplify the PSAM^4^ -GlyR, the plasmid 119739 (Addgene) was used as a template, and the same PCR protocol was followed. However, the annealing gradient was between 65°C and 72°C instead of 55-60°C. In this reaction, the forward primer was (<u>CCGTCAGATCCAAGTTTGTA</u>GGGCTAGCGCCACCATGCGCT), and the reverse primer was (<u>TGTACAAGAAAGCTGGGTCT</u>CCGCTTACTGGTTGTGTACGTC). All PCR reactions were carried out using the KAPA-HiFi enzyme (Roche). The reaction products were run on a gel (1% agarose in TAE). Then, the desired bands were extracted: 633 bp for the first reaction and 1354 bp for the second reaction. The gel extraction was carried out using the Qiagen gel extraction kit following the manufacturer’s protocol.

The in-fusion reaction using the HD in-fusion kit (Takara Bio) was carried out using 100ng of the cut vector, 33.3 ng of the CMV fragment, and 72 ng of the PSAM^4^-GlyR fragment. 1.5µl of the reaction mixture was transformed into Stbl3 cells (Thermofisher) (40 secs at 42C° followed by 2 mins in ice). The recovery of Stbl3 cells was achieved by adding 200µl of SOC medium, and the bacterial cells were incubated at 37°C for 1 hour. The suspension was then poured onto an LB agar (with ampicillin 100µg/ml) plate and was incubated at 37°C overnight. After 16 hours, several colonies were present, and one of them was picked and grown in 5ml of LB ampicillin broth for 6 hours. Then, the 5ml medium was transferred into a 200ml LB ampicillin broth flask and left shaking at 220 RPM at 37°C overnight. Following that, the DNA was extracted using the Qiagen Maxiprep kit following the manufacturer’s protocol. The resulting DNA was verified by sequencing (Source Bioscience). The plasmid was named CMV PSAM^4^ GlyR WPRE BGH pA. To add a sequence encoding for the fluorescent protein mCherry, the CMV PSAM^4^ GlyR WPRE BGH pA plasmid was subjected to an overnight restriction digestion reaction using BsaI-HF v2 (NEB). The restriction reaction products were run on a gel, and the linearized vector (5686 bp) was extracted as described above. A gene fragment corresponding to IRES mCherry was synthesized by Integrated DNA Technologies (IDT), and appropriate 15 bp overhangs on each side were also incorporated in the fragment to allow successful cloning into the linearized CMV PSAM^4^ -GlyR WPRE BGH pA plasmid. 100 ng of cut vector were used with a 3:1 insert to vector molar ratio with IRES mCherry to generate the final plasmid CMV PSAM^4^ GlyR IRES mCherry WPRE BGH pA (see Figure 1) using the in-fusion reaction technique. IRES stands for Internal ribosome entry site, and it is used to allow the translation of an independent protein sequence encoded in the same plasmid without the need to add another promoter. Plasmid CMV PSAM^4^ GlyR IRES mCherry WPRE BGH pA was then tested using *in vitro* GCaMP imaging to assess whether the application of varenicline to PSAM^4^ -GlyR+ DRG neurons could silence the veratridine-evoked calcium responses (see Figure 3).

### Animal work

#### Declaration of animal work

All experiments were performed with the approval of personal and project licenses from the United Kingdom Home Office according to guidelines set by the Animals (Scientific Procedures) Act 1986 Amendment Regulations 2012 and guidelines of the Committee for Research and Ethical Issues of IASP. All experiments involving live animals were approved by the ethical review committee at University College London (UCL). Experiments were carried out blind by experimenters.

#### Compound administration and doses

For behavioral experiments, mice expressing PSAM^4^ -GlyR were intraperitoneally treated with varenicline at a dose of 0.3 mg/kg (in a volume of 10µl/g) or PBS (10µl/g). For experiments involving the intrathecal administration of varenicline, varenicline was injected at a dose of 180ng (in 6µl of PBS).

In preparation for *in vivo* calcium imaging, mice were anesthetized by injecting them intramuscularly with a combination of ketamine (100 mg/kg), xylazine (15 mg/kg), and acepromazine (2.5 mg/kg).

#### Viral injection

All the mice used in experiments involving AAV9 to express PSAM^4^ -GlyR expressed GCaMP3 under the control of the *Pirt* promoter (a kind gift from Prof. Xinzhong Dong (Kim et al., 2014)). Mouse pups (P4-P5) were placed on ice for anesthesia and then injected intraplantarly with a virus packaged by VectorBuilder (CMV PSAM^4^GlyR IRES mCherry WPRE BGH pA) using a 10µl Hamilton syringe with a cannula fused to a 30-G needle. Prior to the injection, the mother was anesthetized using a low concentration of isoflurane (1%). The volume injected in each hind paw was 5µl (1x10^13^ genome copies/ml). After the injection, and until their body temperature returned to normal, pups were maintained in a heating box. Pups were returned to the cage prior to the return of the mother. Mice injected through this route were used for *in vivo* calcium imaging and behavioral tests after they reached adulthood.

#### Transgenic mice and their breeding strategies and genotyping

All transgenic mice used in this study had a C57BL/6 background, and they were housed in groups of 2–5 per cage with a 12-hour light/dark cycle and were allowed free access to water and a standard diet. While we always aimed to have equal numbers of male and female mice, the complexity of transgenic crosses hindered this on some occasions. As our studies were not intended to look for sex differences, both sexes were combined for analysis. All experiments were conducted on adult (more than 6 weeks old) rodents. The individual figure legends for each dataset provide information about the total number of animals used to produce it. For all genotyping experiments, genomic DNA was extracted from ear tissue and then subjected to PCR for testing.

##### Pirt GCaMP3

The primers for genotyping were TCCCCTCTACTGAGAGCCAG (WT forward), GGCCCTATCATCCTGAGCAC (WT reverse), TCCCCTCTACTGAGAGCCAG (GCaMP3 forward) and ATAGCTCTGACTGCGTGACC (GCaMP3 reverse). The *Pirt*-wildtype band size is 300bp and the *Pirt*-GCaMP3 band is 400bp.

##### Na_V_1.8 Cre

The primers for genotyping were CAGTGGTCAGGCTGTCACCA (WT/Cre forward), ACAGGCCTTCAAGTCCAACTG (WT reverse) and AAATGTTGCTGGATAGTTTTTACTGCC (Cre reverse). The Na_V_1.8-wildtype mice band size is 258bp and the Na_V_1.8 Cre band is 346bp.

##### ROSA26-FLEx-PSAM^4^ -GlyR-IRES-tdTomato knockin mice

The ROSA26-FLEx-PSAM^4^ -GlyR-IRES-tdTomato knockin mice were generated using a CRISPR/Cas9 system by Taconic/Cyagen. Briefly, the guide RNA (5’-CTCCAGTCTTTCTAGAAGATGGG-3’) matching the forward strand of the mouse ROSA26 gene, the donor vector (containing “CAG promoter-loxP-lox2272-rBG pA-WPRE-tdTomato-IRES-Kozak-PSAM^4^ -GlyR-loxP-lox2272” cassette) and Cas9 mRNA were co-injected into fertilized mouse eggs to generate targeted conditional knockin offspring. F_0_ founder animals were screened by PCR, followed by sequencing analysis. F_0_ mice were then bred to C57BL/6 mice to test germline transmission. To detect whether germline transmission occurred, the primers (5’-CACTTGCTCTCCCAAAGTCGCTC-3’) and (5’-ATACTCCGAGGCGGATCACAA-3’) were used. The wild-type band was 453 bp and the mutant band was 218 bp.

Upon crossing a Rosa26 FLExed PSAM^4^ -GlyR IRES tdTomato mouse with a homozygous mouse that expresses Cre recombinase in a subset of neurons, some of the resulting offspring will express PSAM^4^ - GlyR and the fluorescent protein tdTomato in that specific neuronal subset. Here, we crossed heterozygous Rosa26 FLEx PSAM^4^ -GlyR IRES tdTomato mice with homozygous Na_V_1.8 Cre mice to obtain experimental animals. For genotyping, three primers were used, TCCACTGCAAGTAGTGATCGGTCC, GGCAACGTGCTGGTTATTGTG, and GCATCTGACTTCTGGCTAATAAAG. When the mouse has a FLEx PSAM^4^ - GlyR IRES tdTomato (i.e., no Cre activity) allele, a 537bp band was seen. When there was Cre activity (i.e., PSAM^4^ -GlyR and tdTomato are produced), one band with either 358bp or 356bp size was seen. Figure 2 shows the breeding strategy for this mouse line.

**Figure 2:**
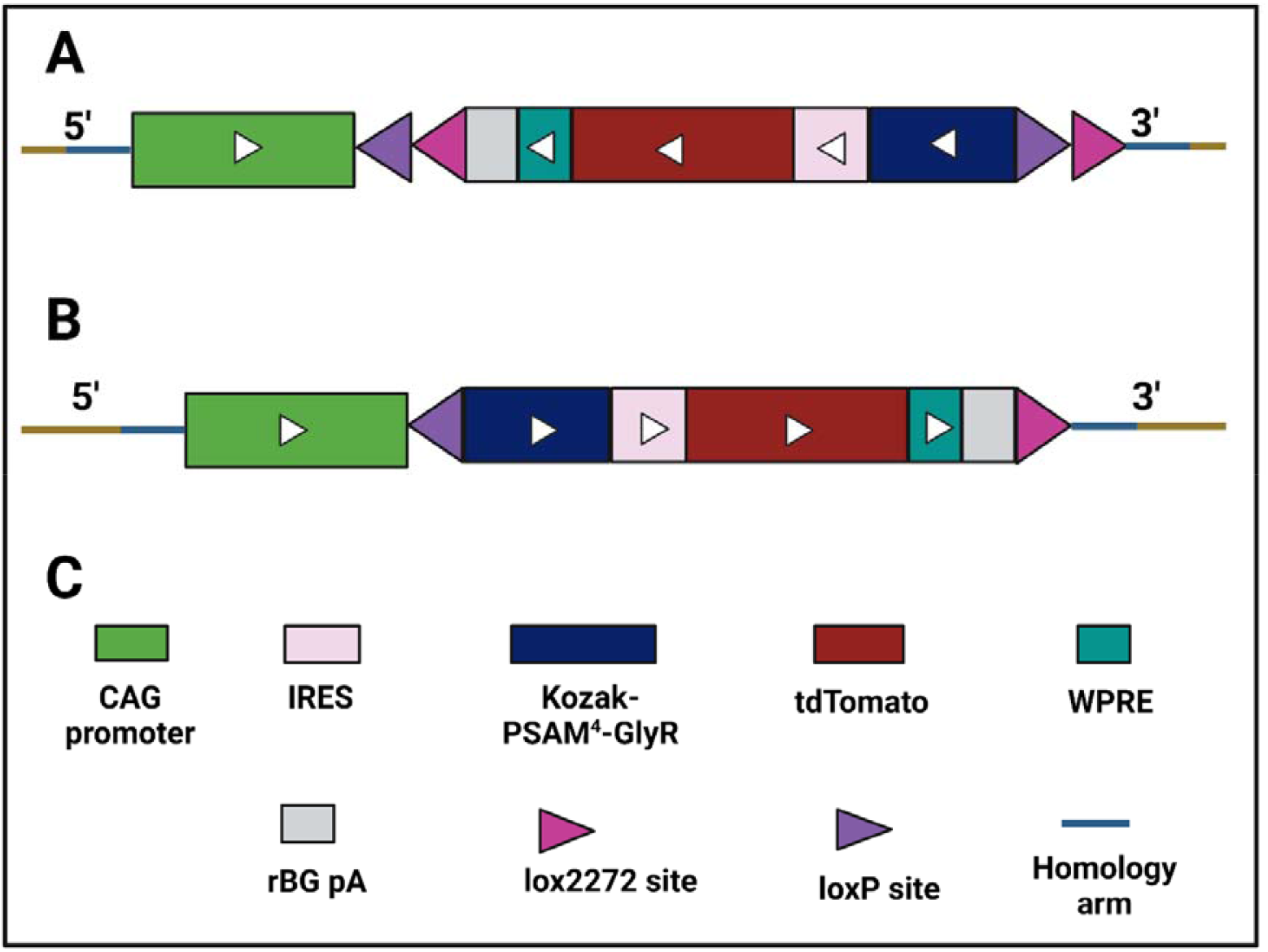
The breeding strategy to produce Rosa26 FLEx PSAM^4^ -GlyR IRES tdTomato and the products after Cre-based recombination. A) shows the mutant allele. B) the product after Cre recombination (PSAM^4^ -GlyR and tdTomato will be produced), and C) shows what each color means.

**Figure 3:**
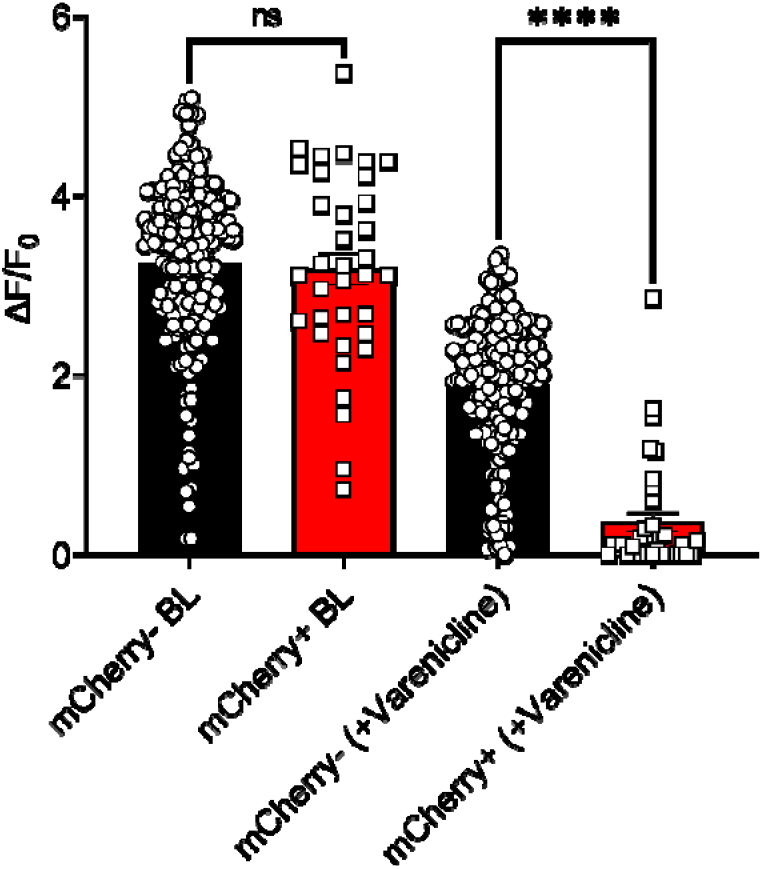
The application of varenicline silenced veratridine-evoked calcium responses in PSAM^4^ -GlyR+ DRG neurons in vitro. Veratridine (30µM) was applied for 10 seconds to measure baseline responses. Varenicline (20nM) was then applied for 5 minutes followed by the application of veratridine (30µM) + varenicline (20nM) for 10 seconds. The comparison of veratridine-evoked increases in calcium signals revealed that PSAM^4^ -GlyR+ DRG neurons (mCherry+, red column) had a similar median response intensity to non-transfected cells (mCherry-, black column) before the exposure to varenicline (p-value= 0.6760, Mann-Whitney test). On the other hand, mCherry+ cells responded significantly less than the PSAM^4^ -GlyR-DRG neurons (mCherry-, black column) after the exposure to varenicline (p-value<0.0001, Mann-Whitney test). n=190 for the mCherry-cells and 35 for the mCherry+ cells. Error bars represent the standard error of the mean. Data were acquired from four independent cultures.

#### Rosa-floxed stop-Diphtheria toxin subunit A (DTA)

To ablate the Na_V_1.8-expressing neurons, heterozygous Rosa-floxed stop-DTA mice (C57BL/6 background) were bred with homozygous mice that express Cre in the Na_V_1.8-positive neurons (C57BL/6 background) (Abrahamsen et al., 2008a).

To check for the presence of a wild-type band in the Rosa-floxed stop-DTA mouse line, the forward primer was AAAGTCGCTCTGAGTTGTTAT, and the reverse primer was GGAGCGGGAGAAATGGATATG. The resulting wild-type band was 600bp in size. To check for the presence of a floxed-stop DTA band in the Rosa-floxed stop-DTA mouse line, the forward primer was AAAGTCGCTCTGAGTTGTTAT, and the reverse primer was GCGAAGAGTTTGTCCTCAACC. The resulting band was 250bp in size.

### Pain models

#### Inflammatory pain

Mice were injected with 3.52µg of prostaglandin E2 (PGE2) intraplantarly (20 µl of a 500 µM solution of PGE2 in saline). Ten minutes after the injection, mice were treated with either varenicline (0.3mg/kg) or PBS intraperitoneally. After 30 minutes of the varenicline/PBS treatment, mice were subjected to the Hargreaves’ test.

#### Chemotherapy-induced cold allodynia

Mice were injected with 80µg (dissolved in 5% glucose solution and 40µl were injected) of oxaliplatin intraplantarly as previously described by (MacDonald et al., 2021a), and after approximately 4 hours, mice were treated with either varenicline or PBS intraperitoneally. 30 minutes after varenicline/ PBS administration, mice underwent the dry ice test.

#### Cancer-induced bone pain (CIBP)

### Cell culture

Lewis lung carcinoma (LL/2) cells (from American Type Culture Collection (ATCC)) were cultured in a medium containing 90% Dulbecco’s Modified Eagle Medium (DMEM) and 10% fetal bovine serum (FBS) and 0.1% Penicillin/Streptomycin for 14 days before the surgery. DMEM was supplemented with L-glutamine (1%) and glucose (4.5 g/liter). The cells were sub-cultured whenever ∼80% confluency was reached, which was done a day before the surgery. On the surgery day, LL/2 cells were harvested by scraping and were centrifuged at a speed of 1500 rpm for two mins. The supernatant was removed, and the cells were resuspended in a culture medium that contained DMEM to attain a final concentration of ∼2x10^6^ cells/ml. The cell counting and viability check were done using the Countess automated cell counter (Thermo Fisher Scientific).

### Surgery

Surgery was carried out on anesthetized mice. Anesthesia was achieved using 1.5-2% isoflurane. The legs and the thighs of the mice were shaved, and the shaved area was cleaned using hibiscrub solution. A sterile lacri-lube was applied to the eyes. The reflexes of the mice to pinches were checked to ensure successful anesthesia. An incision was made in the skin above and lateral to the patella on the left leg. The patella and the lateral retinaculum tendons were loosened to move the patella to the side and expose the distal femoral epiphysis. A 30G needle was used to drill a hole through the femur to permit access to the intramedullary space of the femur. The 30G needle was removed, and a 0.3ml insulin syringe was used to inoculate ∼2x10^4^ LL/2 cells suspended in 10µl of DMEM. The hole in the distal femur was sealed using bone wax (Johnson & Johnson). To ensure that there was no bleeding, the wound was washed with sterile normal saline. Following that, the patella was placed back into its original location, and the skin was sutured using 5–0 absorbable vicryl rapid (Ethicon). Lidocaine was applied at the surgery site, and the animals were placed in the recovery chamber and monitored until they recovered.

In all the studies, male and female C57BL/6 mice (or transgenic mice in C57BL/6 background) were used. The choice of the animal number was based on previous experience from animal experiments in our lab and other labs.

#### Chronic constriction injury of the sciatic nerve

A longitudinal skin incision was made at the level of the right femur after the right thigh region had been shaved and the skin had been cleaned with hibiscrub. The muscular fibers were separated with forceps to make it possible to visualize the sciatic nerve. Following that, three 5-0 silk sutures were loosely tied around the sciatic nerve’s diameter to enable the generation of nerve injury. The skin was sutured to close the surgery site using a 6-0 Vicril suture. Importantly, surgical operations were only carried out when mice were fully anesthetized using isoflurane (1.5–2%).

Following that, lidocaine was applied topically at the surgery site. Animals were then kept in a warm area until fully recovered. Behavior analyses were conducted during the second and third weeks after the operation. This model was shown to induce mechanical, heat, and cold hypersensitivity (Minett et al., 2014; Labuz et al., 2016). For the sham-operated mice, the same steps were followed except for the sciatic nerve ligation.

##### Behavioral tests

All animal experiments were conducted according to Home Office guidelines. The researcher was blind to the treatment. Animals were trained to be handled, and attempts were made to minimize their discomfort during testing. In all experiments, male and female mice were utilized, and the genders and numbers are specified in the figure legends.

##### Hargreaves’ test

The Hargreaves’ test was used to measure acute heat pain thresholds and cutaneous heat hyperalgesia and was performed as previously described by (Hargreaves et al., 1988).

##### Dry ice test

The dry ice test protocol can be found in (Brenner et al., 2012). For this test, dry ice was crushed with a hammer. A 2.5 mL syringe had its top cut off to create the probe’s shape. The modified syringe was filled with dry ice powder, and the open end was placed against a smooth surface while pressure was applied to the plunger to flatten the powder into a dense, 1 cm-in-diameter pellet. To evaluate the withdrawal threshold, the end of the syringe was pressed to the glass beneath the hind paw. The distal joints were avoided, and the center of the right hind paw was targeted, making sure that the paw itself was truly touching the glass surface. A stopwatch was used to measure the time until the withdrawal. The cut-off point for this point was 30 seconds.

##### von Frey test

The “up-down” von Frey was carried out as described by (Deuis et al., 2017). In this test, cutaneous mechanical sensitivity in the hind paw was assessed by evaluating a 50% withdrawal threshold. Mice were habituated for two hours in darkened enclosures with a wire mesh floor. At the beginning of the test, a von Frey filament weighing 0.4g was applied for three seconds. In the subsequent trial, a weaker filament was applied if the response was positive, and a stronger filament was applied if the response was negative. To estimate the 50% withdrawal threshold, five filament application trials were done following the first switch in response (from no response to a positive response or vice versa). The formula utilized to estimate the 50% withdrawal threshold was (10[χ+κδ])/10,000). In this formula, χ represents the log of the last von Frey filament used, δ represents the average variation between filaments used in log units, and κ is a tabular value for the pattern of the responses. The filament weighing 2 grams was used as a cut-off point for this test.

##### Rotarod

To test motor abilities, the rotarod test was carried out as previously described by (Stirling et al., 2005).

##### Acetone evaporative cooling assay

Mice were positioned in plastic chambers on a metal mesh platform. A drop (about 20 µl) of acetone was applied to the plantar surface of the ipsilateral hind paw. After the acetone application, and for 1 minute, the duration of nocifensive behavior was counted in seconds. Nocifensive behaviors included licking, flinching, and lifting. The measurements were repeated three times with a 10-minute gap to determine a mean value for the duration of nocifensive behavior.

##### Weight-bearing

The scale used for this behavioral test was the Incapacitance Metre (Linton Instrumentation), which has two scales to assess the weight put on each hindlimb. The weight placed on each hindlimb is measured for 3 seconds. The readings for the weight put on the hindlimbs were recorded three times for each mouse, and between the readings, mice were allowed to re-place themselves into the tube. The fraction of the weight put on the ipsilateral paw was determined by the summation of all three readings of the weight put on the ipsilateral paw divided by the summation of all the weight measurements on both paws.

##### Randall-Selitto test

The Randall-Selitto test for noxious mechanical sensation was done by applying mechanical pressure on the tail as previously described by (Abrahamsen et al., 2008b; MacDonald et al., 2021b).

### Calcium imaging

#### *In vitro* calcium imaging

##### DRG culture

For each experiment, an adult C57BL/6 mouse expressing GCaMP3 calcium sensor under the control of the *Pirt* promoter was killed by inhalation of a rising CO_2_ concentration followed by cervical dislocation to confirm death. DRGs were dissected from the entire length of the spinal column. During dissection, DRGs were placed in a 35 mm^3^ dish containing Hanks’ balanced salt solution (HBSS) buffer, and the dish was always placed on ice. After the dissection, DRGs were digested in a pre-equilibrated enzyme mix for 45 minutes at 37°C with 5% CO_2_. The composition of the enzyme mix was an HBSS solution containing collagenase (type XI; 5 mg/ml), dispase (10 mg/ml), HEPES (5 mM), and glucose (10 mM). DRGs were then gently centrifuged for 5 minutes at 300 revolutions per minute; the supernatant was discarded and replaced with 1 ml of warmed Dulbecco’s modified Eagle’s medium (DMEM), supplemented with L-glutamine (1%), glucose (4.5 g/liter), sodium pyruvate (110 mg/liter) and 10% FBS. Then, DRGs were triturated mechanically using three fire-polished glass Pasteur pipettes of progressively decreasing inner diameter. The triturated DRGs were then centrifuged at 300 revolutions per minute for eight minutes, the supernatant was discarded, and the cells were resuspended in the needed volume of DMEM, supplemented with L-glutamine (1%), glucose (4.5 g/liter), sodium pyruvate (110 mg/liter), nerve growth factor (50 ng/ml) and 10% FBS. When plasmids were used, the cells were resuspended in P3 buffer from the Amaxa P3 primary cell 4D-Nucleofector X kit, and the plasmid of interest was added to the mixture. Following that, the electroporation protocol was followed according to the manufacturer’s recommended protocol. After the electroporation, a warmed RPMI medium (Thermofisher) was added to the cells, and the cells were left at room temperature for 5 minutes. The DRGs were then centrifuged for 5 minutes at 300 revolutions per minute. To remove the excess plasmid and the excess P3 solution, the supernatant was discarded, and the cells were resuspended in the needed volume of DMEM, supplemented with L-glutamine (1%), glucose (4.5 g/liter), sodium pyruvate (110 mg/liter), nerve growth factor (50 ng/ml) and 10% FBS. Finally, cells were plated onto 9 mm glass coverslips coated with poly-L-lysine (1 mg/ml) and laminin (1 mg/ml). Cells were incubated at 37°C in 5% CO_2_ and calcium imaging experiments were carried out after 48-72 hours post-dissociation and electroporation. When recombinant AAV9 (CMV PSAM^4^ GlyR IRES mCherry WPRE BGH pA, VectorBuilder) was used, cells were plated onto coated 9mm^3^ glass coverslips after dissociation (4 coverslips per 35 mm^3^ dish). 5µl of the virus (1x10^13^ genome copies/ml) was added to the dish. After that, cells were incubated at 37°C in 5% CO_2_, and calcium imaging experiments were carried out after 72 hours post-dissociation.

##### Data Acquisition

The imaging protocol involved applying artificial cerebrospinal fluid (ACSF) for 30 seconds, veratridine (30µM) for 10 seconds, followed by ACSF for 30 seconds, then varenicline (20nM) was applied for 5 minutes, and then a solution containing veratridine (30µM) and varenicline (20nM) was applied for 10 seconds to reassess the response to veratridine. All compounds were dissolved in the ACSF solution. The composition of the ACSF solution was 111 mM sodium chloride, 3.09 mM potassium chloride, 10.99 D-glucose, 25 sodium bicarbonate, 1.26mM magnesium sulfate, 2.52 calcium chloride, and 1.10 potassium phosphate dibasic. Calcium responses to veratridine have been previously characterized by (Mohammed et al., 2017). According to their results, veratridine (30 µM) elicited robust responses in roughly 70% of sensory neurons from C57BL/6 mice. We, therefore, selected this concentration for our experiments.

Using a CCD camera, images were acquired. A 488 nm laser line was used to activate GCaMP3 (1–10% of maximal laser power). A 552 nm laser line operating at 1–15% of maximal laser power was used to stimulate mCherry. The emitted light’s filtering and collecting were adjusted to increase yield and reduce cross-talk (Leica Dye 164 Finer, LasX software, Leica). A hybrid detector with 100% gain was used to find GCaMP and a photomultiplier tube was used to find mCherry (500-600V gain). Images were recorded at a frame rate of 2.81 Hz and a bidirectional scan speed of 800 Hz.

### *In vivo* calcium imaging

#### DRG exposure

Ketamine (100 mg/kg), xylazine (15 mg/kg), and acepromazine (2.5 mg/kg) were used to anesthetize adult mice expressing GCaMP3 (males and females (10-12 weeks old)). Intramuscular injections of anesthetics were administered into the hindlimb on the opposite side of the DRG utilized for imaging. Every 20 to 30 minutes, the animal received a new dosage. The pedal response in each of the four paws was watched in order to gauge the depth of anesthesia. Additionally, further signs of profound anesthesia, like absent whisker movement and a constant respiratory rate, were observed. Only once the pedal reflex was completely gone in all four limbs, whisker movement was gone, and respiration was calm, the procedure was started. Using a heated pad, animals were kept at a constant body temperature of 37°C (VetTech). All surgical equipment was heat-sterilized using a bead sterilizer. At spinal level L3-5, a lateral laminectomy was carried out. In order to expose the spinal column, the skin was longitudinally incised. Using microdissection scissors and an OmniDrill 35, the transverse vertebra and superior articular processes were eliminated. Using microdissection forceps, the dura mater and arachnoid membranes were meticulously dissected to reveal the sensory neuron cell bodies in the ipsilateral DRG. The vertebral column (L1), rostral to the laminectomy, was clamped using a specially constructed clamp to hold the animal in place. Sterilized orthodontic sponges were used to stop bleeding. To reduce breathing-related disturbance, the animal’s trunk was slightly raised. To preserve tissue integrity during the process, artificial cerebrospinal fluid [120 mM NaCl, 3 mM KCl, 1.1 mM CaCl_2_, 10 mM glucose, 0.6 mM NaH_2_PO_4_, 0.8 mM MgSO_4_, 1.8 mM NaHCO_3_ (pH 7.4 with NaOH)] was perfused over the exposed DRG or the DRG was separated by covering with silicone elastomer.

#### Image acquisition

Images were acquired using a Leica SP8 confocal microscope. We utilized a 10x dry, 0.4-NA objective with a 2.2 mm working distance and a 0.75–3x optical zoom to magnify the images. A 488 nm laser line was used to ignite GCaMP3 (1–10% of maximal laser power). A 552 nm laser line operating at 1–15% of maximal laser power was used to stimulate mCherry. The emitted light’s filtering and collecting were adjusted to increase yield and reduce cross-talk (Leica Dye 164 Finer, LasX software, Leica). A hybrid detector with 100% gain was used to find GCaMP3 and a photomultiplier tube was used to find mCherry (500-600V gain). At a frame rate of 1.55 Hz, a bidirectional scan speed of 800 Hz, and a pixel dwell time of 2.44 µs, 512 × 512-pixel pictures were recorded. Using a Pasteur pipette, the left hind paw (ipsilateral to the exposed DRG) was subjected to water heated to 55°C.

### Analysis

The Leica LasX software was used to quantify the calcium signal over time, as indicated by the mean pixel intensity over time. Excessive Z movement in stacks was disregarded for analysis. Stacks from DRGs with abnormal blood flow or surgical injury were also disregarded. Using the free hand tool, regions of interest (ROI) were manually formed all around cells that appeared to be reacting. ROIs covered the whole cell, not just the cytoplasm. A variety of characteristics, including a distinctive cell shape, a visible darker nucleus with less GCaMP signal, an increase in fluorescence after sensory stimulation, the lack of fluctuating fluorescence changes not tied to the stimulus, suggestive of non-sensory activity, and the absence of a continuous high GCaMP signal, indicative of cell death or injury, were necessary for manually identifying responsive cells. The mean pixel intensity for the antecedent 15 seconds prior to stimulus application was computed. Neurons were classified as responders if the maximum derivative was greater than the baseline derivative plus four standard deviations (Z-score ≥ 4). Then, for each response, the ∆F/F_0_ ratio was calculated to derive a normalized quantitative measure of the fluorescence alteration. Subsequently, the neurons that were deemed responsive were analyzed statistically.

### DRG sections preparation and staining

Mice were killed using a rising CO_2_ concentration and death was confirmed by cervical dislocation. Following that, the lumbar DRGs (L2-L4) were extracted and placed in 4% paraformaldehyde (Sigma-Aldrich) dissolved in 0.1M phosphate-buffered saline (PBS) (pH = 7.4) (overnight at 4 °C). The solution was then removed, and the DRGs were cryoprotected overnight in 30% w/v sucrose in 0.1M PBS at 4°C. The DRGs were embedded in molds with OCT embedding compound (Tissue-Tek®, Sakura) and left to set on dry ice, then stored at -80°C till sectioning. The whole DRG was sectioned at 11µm thickness and was collected on electrostatically charged slides (Superfrost® Plus, Thermo-Scientific). Slides were permitted to dry at room temperature. The sections were then washed using 1x PBS (three times, for five minutes each at room temperature). Then the sections were permeabilized using 1% Tween in 1xPBS (for 1 hour at room temperature). Following that, the sections were washed using 1xPBS for 5 minutes at room temperature. The non-specific binding was blocked by the application of a blocking buffer at 4°C overnight. The blocking buffer was prepared using 5% w/v milk powder (Sigma) in 1% tween 20 in 1xPBS. Then the sections were incubated in the diluted primary antibody solution dissolved in blocking buffer (overnight at 4°C). The sections were then washed using 0.1% tween 20 in 1x PBS (three times, 10 minutes each, at room temperature). The sections were then incubated in a solution of the blocking buffer that contained the secondary antibody. The slides were kept in the dark (overnight at 4°C). The slides were then washed three times using 0.1% tween 20 in 1x PBS (5 minutes each at room temperature). They were allowed to dry before the mounting medium was added. Images were acquired using a Leica confocal microscope.

To image tdTomato, rabbit anti-red fluorescent protein (anti-RFP) antibody was used (Rockland, 600-401-379, 1:500), and the secondary antibody was Alexa Fluor 647 anti-rabbit (Jackson ImmunoResearch, 711-605-152, 1:500). To image peripherin, mouse anti-peripherin antibody was used (Sigmdrich, P5117, 1:8000), and the secondary antibody was goat anti-mouse IgG conjugated Alexa 488 (life technologies, A11001, 1:1000).

### Statistical analyses

The mixed-effects model (RMEL) or the two-way analysis of variance (ANOVA) was used with multiple comparisons to compare the pain-like behavior between two groups over time. The RMEL was used when data points were missing, followed by post hoc analyses. In all statistical tests, the difference between groups is considered significant when the p-value is <0.05. All statistical analyses were performed using GraphPad Prism 9. Data are presented as mean ± standard error of the mean (SEM). To compare any two groups, the t-test was performed when the samples passed the D’Agostino & Pearson normality test. If one of the two groups did not pass the normality test, the Wilcoxon test was used for paired comparisons, while the Mann-Whitney test was used for unpaired comparisons. The one-way ANOVA test was used to compare three or more groups at a single time point, followed by Tukey’s test. Statistical significance is expressed as: *p < .05; **p < .01; ***p < .001, ****p < .0001.

## Results

We first cloned a plasmid encoding PSAM^4^ -GlyR and mCherry under the control of the CMV promoter (see Figure 1). We then tested the ability of activated PSAM^4^ -GlyR to silence DRG sensory neurons *in vitro* using GCaMP imaging. Initially, the plasmid vector was used to enable the expression of PSAM^4^ - GlyR in primary DRG cultures from mice expressing GCaMP3 under the control of the *Pirt* promoter. The application of varenicline (20nM) for 5 minutes silenced the calcium responses evoked by the application of veratridine (30µM) in DRG neurons that express PSAM^4^ -GlyR (Figure 3).

With these promising *in vitro* silencing results, we packaged our cloned plasmid into AAV9 viral vectors for further testing. When used *in vitro*, AAV9 vectors enabled the expression of PSAM^4^-GlyR. Veratridine-evoked calcium responses were silenced after varenicline application in transduced cells only, without affecting calcium responses in non-transduced cells (Figure 4).

**Figure 4:**
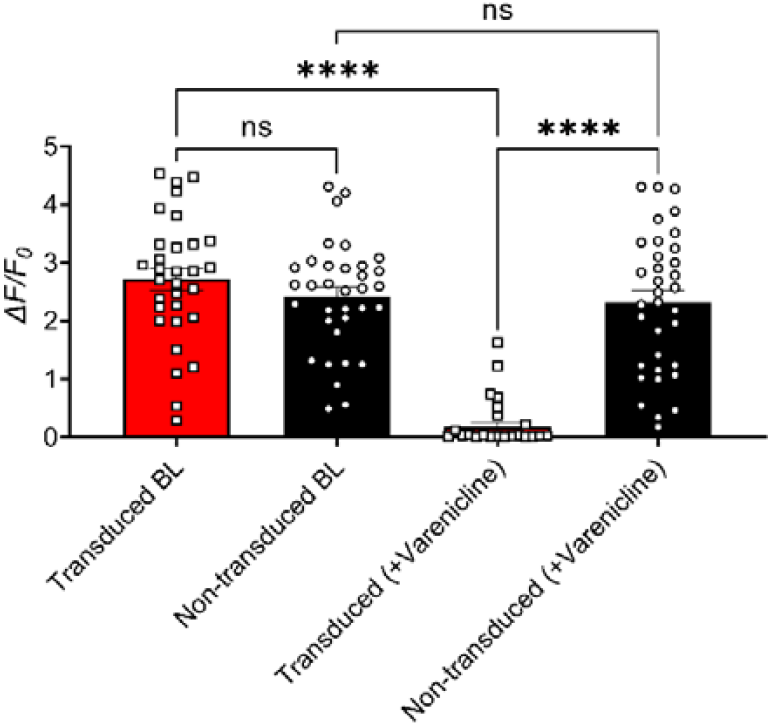
Varenicline (20nM) silenced the responses of PSAM^4^ -GlyR expressing DRG neurons to veratridine (30µM) following recombinant AAV9 vector-based transduction. There was no significant difference in the baseline (BL) response to veratridine (30µM) between transfected and non-transfected cells (unpaired t-test, p-value= 0.2448), but after the exposure to varenicline (20nM) for 5 minutes, the transduced cells responded to veratridine (30µM) significantly less than the non-transduced cells (p-value <0.0001, Mann Whitney test) and significantly less than their responses in the baseline (p-value <0.0001, Wilcoxon matched-pairs signed rank test). There was no significant difference between the responses of the non-transduced cells to veratridine before and after varenicline (p-value= 0.6057, paired t-test). Transduced cells are the DRG neurons expressing PSAM^4^-GlyR as indicated by mCherry positivity. n=31 for the transduced and 34 for the non-transduced.

Following these *in vitro* experiments, AAV9 was tested in live animals expressing GCaMP3 under the control of the *Pirt* promotor to test potential silencing of calcium responses in DRG neurons after varenicline administration. These *in vivo* experiments demonstrated that heat-responding cells were significantly silenced in mice expressing PSAM^4^ -GlyR treated with varenicline (i.p., 0.3 mg/kg) (Figure 5). Because AAV9-based PSAM^4^ -GlyR expression, in conjunction with varenicline, could silence DRG neurons both *in vitro* and *in vivo*, we tested this system in withdrawal thresholds in pain behavioral tests and in inflammatory pain states and chemotherapy-induced cold allodynia (Figure 6), with positive outcomes in pain behavior but also with a reduction in innocuous touch sensation.

**Figure 5:**
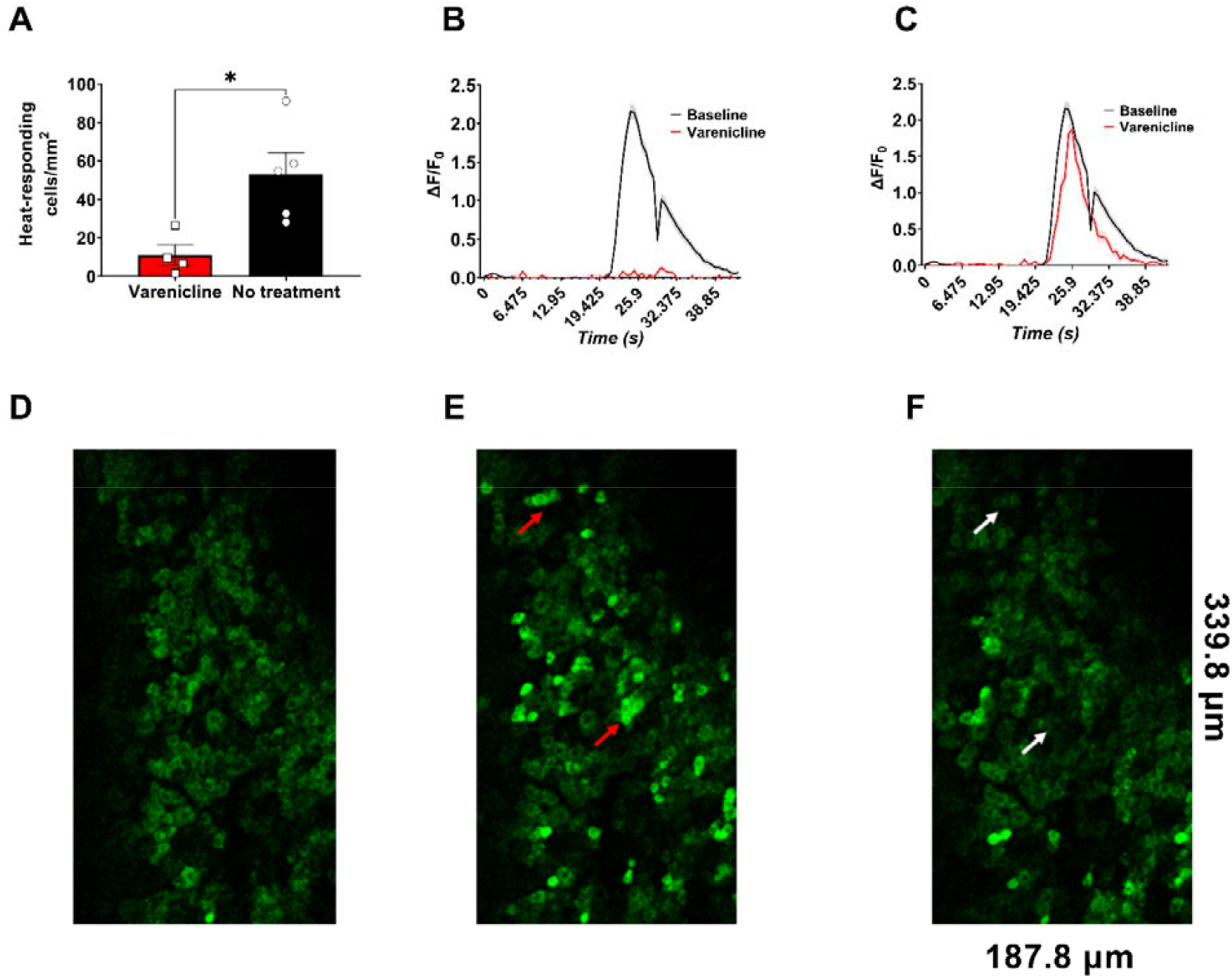
Varenicline reduced the calcium responses of DRG neurons responding to the application of 55°C water to the ipsilateral hind paw of mice expressing the PSAM^4^ -GlyR following recombinant AAV9 vector-based transduction. A) The number of DRG neurons per mm^2^ (in L4) responding to hot water in the paw was reduced significantly after the intraperitoneal administration of varenicline to PSAM^4^ -GlyR expressing mice (p-value= 0.0175 (un-paired t-test, n=5 in the control group and four in the treatment group)). B) Plot showing the mean response amplitude before (black trace) and after varenicline administration in the silenced neurons (n = 98 cells for the varenicline-treated group and 147 for the baseline). C) Plot showing the mean response amplitude before (black trace) and after varenicline administration in the silencing-resistant neurons (n = 49 cells for the varenicline-treated group and 147 for the baseline). D) represents the basal calcium signal in L4 (no stimulation) in mice expressing GCaMP3 and PSAM^4^ -GlyR. E) shows the increase in the calcium signal in the ipsilateral DRG after the application of 55°C water to the ipsilateral paw (no treatment). Red arrows point towards representative heat-responding cells. F) shows the response of the same DRG to the application of 55°C water to the paw 30 minutes after the mouse received an intraperitoneal injection of varenicline (0.3 mg/kg) (The white arrows point toward the cells that previously responded to heat before varenicline application). Data are presented as mean±SEM.

**Figure 6:**
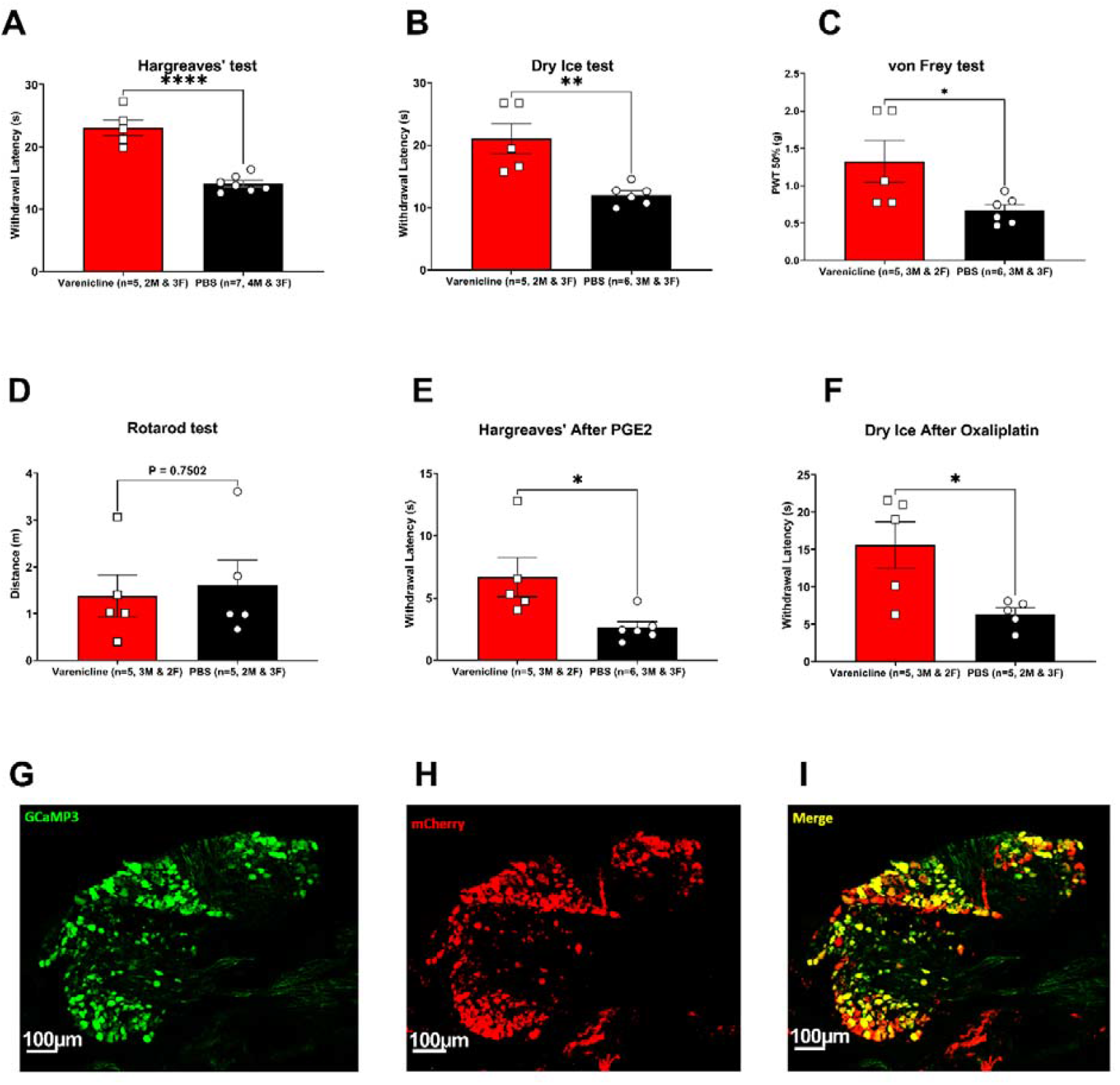
Varenicline application increased the withdrawal thresholds of mice expressing PSAM^4^ -GlyR in sensory tests and reduced inflammatory hyperalgesia and oxaliplatin-induced cold allodynia without impairing motor coordination. A) Hargreaves’ test (p-value= <0.0001). B) von Frey (p-value= 0.0373). C) Dry ice test (p-value= 0.0035). D) Rotarod test (p-value=0.7502). E) 3.52µg of PGE2 was injected intraplantarly, and after 10 minutes, either varenicline or PBS was injected intraperitoneally. After 40 minutes of the PGE2 injection, the Hargreaves’ test was performed. The varenicline-treated mice had significantly less heat hyperalgesia following the intraplantar injection of PGE2 compared to the PBS-treated group (p-value= 0.0249). F) The administration of varenicline to mice expressing PSAM^4^ -GlyR in the DRG neurons abolished the cold allodynia driven by the intraplantar injection of 80µg oxaliplatin (p-value= 0.0209). Figures G-I show the expression of PSAM^4^ -GlyR 10 weeks after injecting a recombinant AAV9 carrying PSAM^4^ -GlyR and mCherry transgenes into mouse pups that express the calcium sensor GCaMP3 under the control of the Pirt promoter. The green color (G) represents DRG neurons expressing GCaMP3; red represents DRG neurons expressing PSAM^4^ -GlyR (H), and Figure (I) shows a merged picture of G and H. Error bars represent SEM. All statistical analyses were done using the unpaired t-test. Varenicline was injected intraperitoneally at 0.3mg/kg dose, and behavioral tests were done approximately 30 minutes after varenicline administration. See Figure (6-1) for the effect of intrathecal varenicline and for varenicline washout.

To enable the expression of PSAM^4^ -GlyR in defined neuronal subsets, we designed a mouse line that expresses PSAM^4^ -GlyR receptors driven by a CAG promoter in a Cre-dependent manner. We then used Na_V_1.8 Cre expressing mice to selectively target these neurons. In this mouse line, we conducted acute pain behavioral tests (Figure 7) and chronic pain models (Figures 8 and 9) and we found a selective reduction in pain without any detectable effects on innocuous touch sensation.

**Figure 7:**
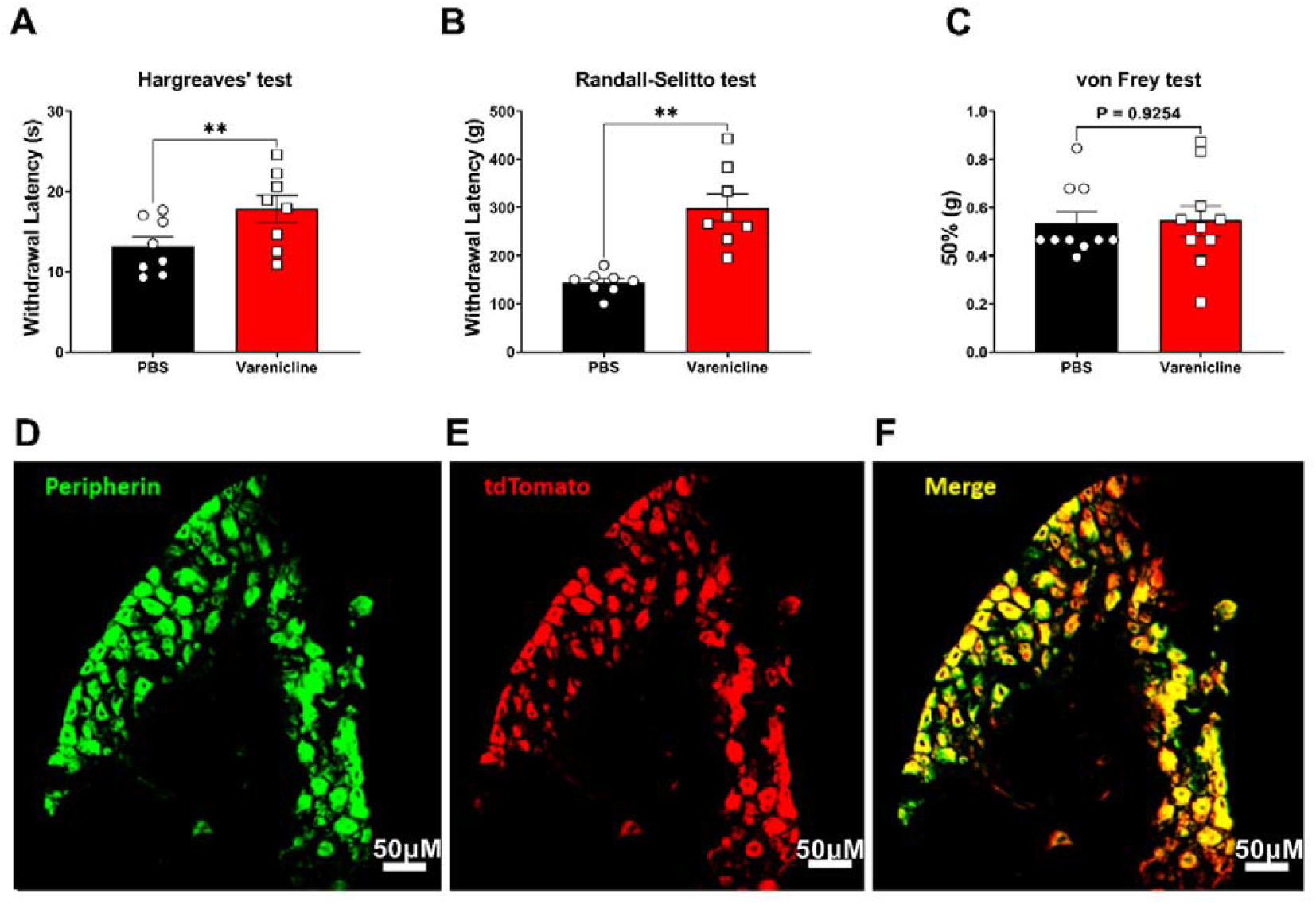
Varenicline treatment to mice that express PSAM^4^ -GlyR in the Na_V_1.8 expressing neurons elevated their withdrawal thresholds in the Randall-Selitto test and the Hargreaves’ test without impairing innocuous touch sensation. Figure (A) shows the Hargreaves’ test results (p-value= 0.0046 paired t-test, n=8 (3M & 5F)), Figure (B) shows the Randall Selitto test results (p-value= 0.0011, paired t-test, n=8 (3M & 5F)), and Figure (C) shows how the systemic administration of varenicline caused no significant alteration in the withdrawal thresholds of the mice expressing PSAM^4^-GlyR in the Na_V_1.8-positive neurons in the von Frey test (p-value=0.9254, paired t-test, n=10 (5M & 5F))). Error bars represent SEM. Varenicline was administered intraperitoneally at a dose of 0.3 mg/kg. Figures (D-F) show the expression of PSAM^4^ -GlyR in the DRGs from mice carrying a Rosa26-FLEx-PSAM^4^ -GlyR-IRES-tdTomato allele and expressing Cre enzyme in the Na_V_1.8-positive neurons. D: peripherin expressing DRG neurons; E: PSAM^4^ -GlyR expressing DRG neurons (as demonstrated by tdTomato expression); F: a merged picture of D and E showing the co-expression of peripherin and PSAM^4^ -GlyR. Images were obtained using a Leica SP8 confocal microscope. Scale bars represent 50µm.

**Figure 8:**
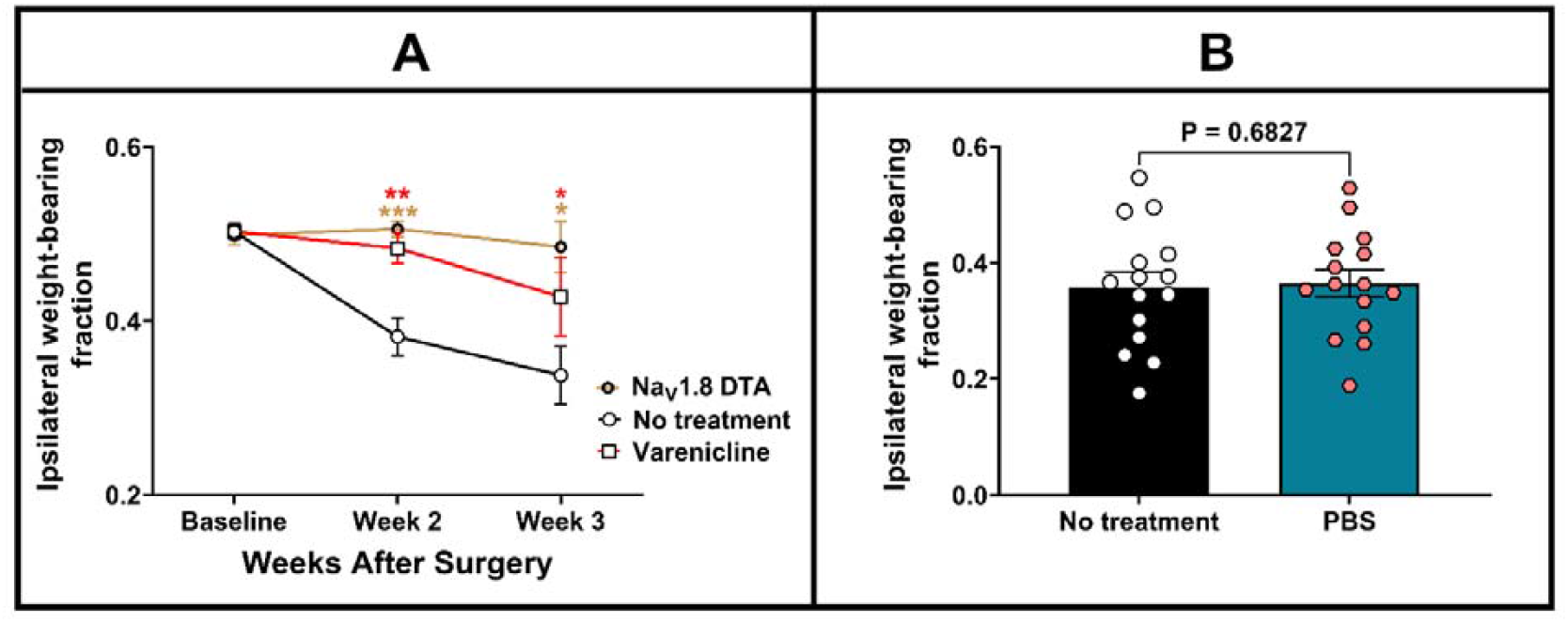
The systemic injection of varenicline in mice expressing PSAM^4^ -GlyR in Na_V_1.8-positive neurons partially reversed weight-bearing deficits caused by CIBP. The weight-bearing test was conducted without any treatment. After that, and on the same day, each mouse received an intraperitoneal injection of either varenicline (0.3 mg/kg) or PBS, and the weight-bearing test was repeated 30 minutes after treatment. Figure (A) compares the weight bearing of three groups of mice (mice in which the Na_V_1.8+ neurons are ablated (Na_V_1.8-DTA, brown line), mice that express PSAM^4^ -GlyR in the Na_V_1.8+ neurons that received no treatment (black line, control) and mice that express PSAM^4^ -GlyR in the Na_V_1.8+ neurons after receiving varenicline (red line)). The comparison of the average weight-bearing fraction on the affected limb between the three groups over time indicated a distinct difference between them, with an apparent reduced pain-like behavior displayed by the Na_V_1.8-DTA group than the control group (the RMEL analysis, p-value <0.0001) and between the varenicline treated group and the control group (RMEL, p-value=0.0026). On the other hand, the difference between the varenicline-treated group and the Na_V_1.8 DTA group is statistically insignificant (RMEL, p-value= 0.3066). The asterisks of significance shown in the figure represent the post hoc analysis results comparing each treatment with the control group at each time point (Sidak’s multiple comparisons test). N=15 (10 females and five males) for the PSAM^4^ -GlyR expressing mice at the baseline and seven (three males and four females) for the Na_V_1.8-DTA group. DTA: Diphtheria toxin subunit A. Figure (B) demonstrates that the administration of PBS to mice that express PSAM^4^ -GlyR in the Na_V_1.8+ neurons with cancer in the femur does not alter their weight-bearing results. Data were analyzed using the paired t-test (p-value shown on the graph). The experiment was conducted on day 13 after the surgery before (black column) or 30 minutes represent the standard error of the mean.

**Figure 9:**
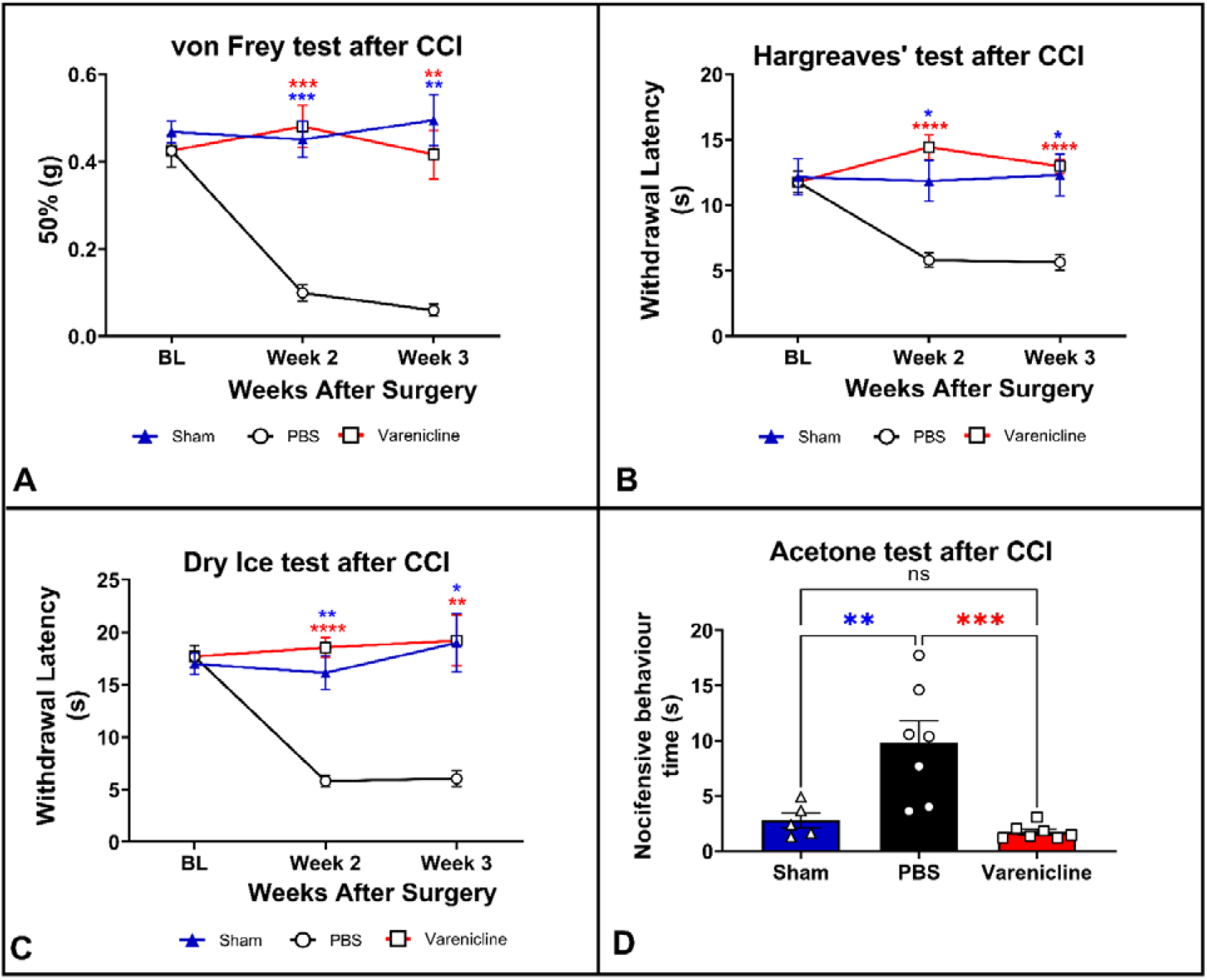
Intraperitoneal administration of varenicline to mice expressing PSAM^4^ -GlyR in the Na_V_1.8+ neurons reversed mechanical, thermal, and cold sensitivity caused by the chronic constriction injury of the right sciatic nerve. Each mouse received either PBS or varenicline and the behavioral tests were carried out 30 minutes after drug administration. 24 hours after that, and within the same week, each mouse received the other treatment, and the behavioral tests were repeated. In graphs (A-D), mice with ligated sciatic nerves (and treated with either varenicline (red) or PBS (black)) are compared with sham mice in which the sciatic nerve was exposed but not ligated (blue). Figure A shows the up-down von Frey test results for mechanical sensitivity. Figure B illustrates the Hargreaves’ test results (heat sensitivity). Figure C shows the dry ice test results (cold sensitivity). Figure D shows the acetone test results on week 3 after the chronic constriction injury of the sciatic (affective responses to cold). For Figures A-C, data were analyzed using the two-way ANOVA with Tukey’s multiple-comparisons test, but for Figure D, the one-way ANOVA test with Tukey’s multiple-comparisons test was employed. n=7 for mice expressing PSAM^4^ -GlyR in the Na_V_1.8+ neurons (three males and four females), and for the sham group, n=5 mice (four males and one female). The varenicline-treated group (red asterisks) or the sham group (blue asterisks) with the PBS-treated group at each time point. Varenicline was used at a dose of (0.3mg/kg) and was injected intraperitoneally. Error bars represent SEM.

### 1 Delivering the PSAM^4^-GlyR transgene using plasmid vectors

We expressed PSAM^4^ -GlyR by electroporating primary DRG cultures from mice that express GCaMP3 in all sensory neurons under the control of the *Pirt* promoter using plasmid vectors. We applied veratridine (30µM) for 10 seconds to measure baseline response, and results indicated that transfected (mCherry+) and non-transfected (mCherry-) DRG neurons had similar responses (Figure 3). Then varenicline (20nm) was applied for 5 minutes, followed by re-activation using veratridine (30µM) in combination with varenicline (20nM) for 10 seconds. In those cells transfected with the PSAM^4^ -GlyR construct, varenicline caused a profound diminution in calcium responses, with the mean calcium response (∆F/F0) being 1.9 in the non-transfected group and 0.37 in the transfected group (Figure 3). Some non-transfected cells which were mCherry negative also showed a small diminution in calcium responses that may reflect the presence of low-level PSAM^4^ -GlyR transduction below the detection level of mCherry fluorescence. With these promising results, the plasmid was packaged into AAV9 vectors to enable further testing.

### 2 Delivering PSAM^4^-GlyR transgene using AAV9

#### *In vitro* experiments

A recombinant AAV9 carrying PSAM^4^ -GlyR and mCherry was used in primary DRG cultures from mice that express GCaMP3. We used the lipid-soluble neurotoxin veratridine (30µm) as a sodium channel agonist to activate these cells. Following that, varenicline (20nm) was applied to test whether the re-application of veratridine after varenicline would cause a similar response in transduced (PSAM^4^ -GlyR expressing) and non-transduced (PSAM^4^ -GlyR negative) DRG neurons. Results from these experiments indicated that varenicline silenced the responses of more than 80% of DRG neurons that express PSAM^4^ - GlyR on veratridine activation and lowered the mean response intensity in the remaining 20% of cells by more than 70%. Varenicline did not significantly affect the veratridine-evoked responses in non-transduced cells (Figure 4). The baseline responses to veratridine were similar between transduced and non-transduced cells, showing that the expression of the receptor is not sufficient to alter the behavior of the cell (Figure 4).

#### *Ex-vivo* viral expression analysis

To assess whether PSAM^4^ -GlyR (+varenicline) system could silence DRG neurons *in vivo*, mouse pups that express GCaMP3 under the control of the *Pirt* promoter were injected intraplantarly with recombinant AAV9 vectors carrying PSAM^4^ -GlyR and mCherry transgenes. When these mice reached adulthood, lumbar DRGs (L2-L4) were extracted and imaged. This experiment was carried out 10 weeks after the intraplantar injections of 5µl (10^13^ genome copies/ml) of recombinant AAV9 in each paw in mouse pups. The imaging revealed a high transduction efficiency and colocalization of PSAM -GlyR expression and GCaMP3 expression, with more than 70% of the GCaMP3+ neurons being mCherry+ (Figure 6 (G-I)). We then moved to test whether the activation of PSAM^4^ -GlyR *in vivo* could also silence DRG neurons *in vivo*.

#### In vivo experiments

For *in vivo* experiments, we used recombinant AAV9 vectors to deliver PSAM^4^ -GlyR and mCherry transgenes. The recombinant virus was injected intraplantarly in mouse pups that express GCaMP3 under the control of the *Pirt* promoter. Mice at 7-16 weeks were used for acute pain behavioral tests, including a model of chemotherapy-induced cold allodynia and a model of inflammation, as well as *in vivo* calcium imaging of the DRG neurons. In these tests, mice were either treated with varenicline or with a control vehicle (PBS). Data consistently showed that varenicline silenced DRG neurons, elevated withdrawal thresholds in behavioral pain tests, and reversed cold and/or heat hypersensitivity after chemotherapy or inflammation (Figure 6).

#### In vivo calcium imaging

*In vivo*, calcium imaging was used to examine whether activating PSAM^4^ -GlyR with varenicline could silence DRG neurons *in vivo* after AAV9-based transgene delivery (Figure 5). The *in vivo* imaging experiments indicated that there was a significant reduction in the number of cells responding to the application of 55°C water to the paw in the ipsilateral L4 DRG neurons 30 minutes after the intraperitoneal administration of varenicline (0.3 mg/kg) to PSAM^4^ -GlyR expressing mice (Figure 5 (A)). A significant reduction in the number of heat-responding cells per mm^2^ in L4 DRG was observed. These results indicate that approximately 80% of the heat-responding cells were silenced in the varenicline-treated group. Figures 5B and 5C show the average peak intensity trace of the cells before the application of varenicline (baseline response) compared to the silenced neurons after varenicline administration (B) or the cells that remained responsive after varenicline (C). For the peak intensity analysis, 147 cells were analyzed, and two-thirds of them were silenced (98 cells) after varenicline administration (shown in Figure 5 B). Figure 5 (D-F) shows examples of representative *in vivo* calcium imaging experiments.

#### Sensory tests and motor coordination

The intraperitoneal injection of varenicline (0.3mg/kg) rendered the withdrawal latency/thresholds of mice expressing PSAM^4^ -GlyR significantly longer/higher than the control group (expressing PSAM^4^ -GlyR but injected with PBS instead of varenicline) in sensory tests. The behavioral tests included the Hargreaves’ test, the von Frey test, and the dry ice test (see Figure 6 (A-C), respectively). These behavioral tests were carried out 30 minutes after the drug injection. In the Hargreaves’ test, the mean withdrawal latency was 23.10 seconds for the varenicline-treated mice, while it was 14.11 seconds for the PBS-treated group (Figure 6 A). Similarly, PSAM^4^ -GlyR activation elevated the withdrawal latency in the dry ice test to become 21.07 in the varenicline group, while the PBS-treated group was 12.03 seconds (Figure 6 B) (p-value <0.0001, unpaired t-test). Importantly, varenicline almost doubled the mean withdrawal threshold in the von Frey test compared to the control group (0.67 grams vs. 1.32 grams), indicating an impairment of innocuous touch sensation (Figure 6 C). To assess whether the administration of varenicline to PSAM^4^-GlyR-expressing mice caused any noticeable impairment in motor abilities, the rotarod test was done, and mice treated with varenicline were compared to mice treated with PBS. No significant difference was detected between the two groups (Figure 6 D).

We assessed whether the expression of AAV9-PSAM^4^ -GlyR alone had any effects on pain behavior. Mice expressing GCaMP3 under the control of the *Pirt* promoter were compared to their littermates that were injected with AAV9 to induce the expression of PSAM^4^ -GlyR in the Hargreaves’ test. Results indicated that the two groups behaved identically (extended figure 6-1B). This excludes the possibility that viral injection alone is altering pain thresholds through an inflammatory mechanism. To confirm that the *in vivo* effects seen with varenicline are caused by its action on PSAM^4^ -GlyR+ DRG neurons, varenicline (or PBS) was administered intrathecally, and the Hargreaves’ test was carried out at various time points. Results demonstrate that intrathecal varenicline also raises the withdrawal threshold of PSAM^4^ -GlyR expressing mice (extended figure 6-1B). We also assessed whether mice could return to their baseline latency in the Hargreaves’ test after washout and results indicated that near-baseline measurements were achieved 6 hours after the intraperitoneal injection of varenicline (0.3 mg/kg) to mice expressing PSAM^4^-GlyR (extended figure 6-1A).

#### Pain models

##### Inflammatory pain

Following initial validation steps, the PSAM^4^-GlyR (+ varenicline) system was tested in a reversible inflammatory model involving the injection of PGE2 intraplantarly. The pro-inflammatory actions of PGE2 involve the activation of protein kinases A and C, which results in heat hyperalgesia (Sachs et al., 2009) as these kinases sensitize many receptors and ion channels, like TRPV1 channels (Kawabata, 2011). The PSAM^4^-GlyR (+varenicline) system showed good efficacy in reducing signs of heat hypersensitivity that usually follow PGE2 injection when tested using the Hargreaves’ test (Figure 6 (E)). While mice treated with varenicline still exhibited signs of heat hyperalgesia, this hyperalgesia was significantly less than the ones treated with PBS, with the mean withdrawal latency in the Hargreaves’ test in the varenicline-treated mice being almost triple that of the PBS-treated group (6.69 seconds vs 2.62 seconds) (unpaired t-test, p-value=0.0249, Figure 6E).

##### Oxaliplatin-induced cold allodynia

PSAM^4^-GlyR expressing mice were injected with the chemotherapeutic agent oxaliplatin intraplantarly. Oxaliplatin is known to induce cold allodynia (demonstrated by the reduction in the withdrawal latency in the dry ice test). This allodynia was reversed in the PSAM^4^-GlyR-expressing mice treated with varenicline compared to the ones treated with PBS (Figure 6F) (unpaired t-test, p-value= 0.0209).

### 3 A Cre-dependent transgenic mouse expressing PSAM^4^-GlyR

Due to the inherent variability in receptor expression when viral vectors are used, we designed a mouse line that expresses PSAM^4^ -GlyR receptors in a Cre-dependent manner to ensure consistency of results and to provide a tool for mechanistic studies. A Rosa26-FLEx-PSAM^4^-GlyR-IRES-tdTomato was generated as described in the methods section. As Na_V_1.8+ neurons are a major nociceptive population in mouse DRG, we used the Na_V_1.8 Cre mouse to unmask PSAM^4^-GlyR expression and examined the effect of silencing this set of neurons on pain behavior.

### Acute and chronic pain models

#### Acute pain models

Mice that express PSAM^4^-GlyR in the Na_v_ 1.8-positive neurons were subjected to various acute pain behavioral tests (the Hargreaves’ test for heat sensation and the Randall-Selitto for noxious mechanical sensation). Results from these tests indicate that intraperitoneal varenicline treatment of mice expressing PSAM^4^-GlyR in Na_v_ 1.8+ neurons doubled their withdrawal thresholds in the Randall-Selitto test (143.75 g after PBS treatment and 299.38 g after varenicline treatment, p-value= 0.0011, paired t-test) and increased the mean withdrawal threshold in the Hargreaves’ test from 13.16 seconds to 17.80 seconds (p-value= 0.0046, paired t-test) (Figure 7A and B). Varenicline (0.3 mg/kg, i.p.) did not significantly impact innocuous mechanical sensation when tested using the up-down von Frey test (Figure 7 (C)), which is considered a main advantage over the use of viruses that do not restrict the expression of PSAM -GlyR to a specific neuronal subtype.

To evaluate the expression of PSAM^4^-GlyR, DRGs from heterozygous Rosa26-FLEx-PSAM^4^-GlyR-IRES-tdTomato mice that express Cre recombinase in the Na_V_1.8-positive neurons were extracted, fixed, and stained for peripherin (a marker for small diameter peripheral neurons (Szabolcs et al., 1996)) and tdTomato. Results are shown in Figure 7 (D-F), and they indicate a high degree of colocalization, with approximately 90% of the peripherin+ neurons showing tdTomato expression.

#### Chronic pain models

##### Cancer-induced bone pain

Mice that express PSAM^4^-GlyR channels in the Na_v_1.8 expressing neurons were subjected to CIBP surgery. Following CIBP, mice show a reduction in the weight placed on the ipsilateral paw, which can be assessed by the incapacitance tester. The ipsilateral weight-bearing fraction was assessed weekly, after the surgery. Following each assessment, mice received an intraperitoneal injection of varenicline (0.3 mg/kg) or PBS, and ipsilateral weight-bearing was reassessed. Varenicline diminished the weight-bearing deficits exhibited by cancer-bearing mice that express PSAM^4^-GlyR in the Na_V_1.8-positive neurons (Figure 8A, red vs. black) (RMEL, p-value 0.0026). Cancer-bearing mice that express PSAM^4^ -GlyR in the Na_V_1.8 positive neurons (and treated with varenicline) were compared with cancer-bearing mice in which the Na_V_1.8-expressing neurons are congenitally ablated (20). The difference in weight-bearing between these two groups was statistically insignificant (Figure 8A, red vs. brown, RMEL, p-value= 0.3066). It is important to note that treating the PSAM^4^ -GlyR expressing cancer-bearing mice with PBS did not change their weight-bearing results significantly (Figure 8B).

##### Neuropathic pain

Mice that express the PSAM^4^ -GlyR channels in the Na_V_1.8-expressing neurons were subjected to chronic constriction injury surgery of the sciatic nerve. After the surgery, mice demonstrated signs of increased mechanical, heat, and cold hypersensitivity (Figure 9 A-D, black). Then these mice received an intraperitoneal injection of varenicline (0.3 mg/kg), and the results showed that varenicline, compared to PBS treatment, reversed signs of mechanical (Figure 9A, red vs. black, two-way ANOVA, p-value=<0.0001), heat (figure 9B, red vs. black, two-way ANOVA, p-value=<0.0001) and cold hypersensitivity ((figure 9C, dry ice test, red vs. black, two-way ANOVA, p-value=<0.0001) (figure 9D, acetone test, red vs. black, one-way ANOVA test with Tukey’s test, p-value=0.0009)).

Injured mice that express PSAM^4^-GlyR in the Na_V_1.8 positive neurons treated with varenicline were also compared with sham mice, and the difference in the von Frey (two-way ANOVA, p-value= 0.4278), Hargreaves’ (two-way ANOVA, p-value= 0.4626), dry ice (two-way ANOVA, p-value= 0.5148) and acetone tests (one-way ANOVA test with Tukey’s test, p-value=0.8554) was statistically insignificant between these two groups in all tests (Figure 9 (A-D), red vs. blue). However, in the absence of varenicline, there was dramatic sensitization of sensory neurons (Figure 9 (A-D), black vs. blue (p-value <0.0001 for the von Frey and the dry ice tests (two-way ANOVA), 0.0064 (for the Hargreaves’ test, two-way ANOVA), and 0.0059 (for the acetone test, one-way ANOVA with Tukey’s test)).

## Discussion

### Genetic tools to silence DRG neurons

Several options exist to silence neurons, including the use of the CRISPR-Cas9 system to delete genes encoding ion channels required for neuronal activation, such as voltage-gated sodium channels. However, CRISPR-Cas9-based gene editing carries some disadvantages, the most important of which is the permanent change in the genome which can occur either at the site of action or at off-target sites (Ledford, 2020). Chemogenetics represents an attractive option that is efficient, significantly less invasive, reversible, and extremely specific. Chemogenetics, when combined with an intersectional approach, can also be applied for cell-type and projection-specific dissection (Lee et al., 2020). Engineered GPCRs used in DREADD and RASSL (Receptors activated solely by selective ligands) (Pei et al., 2008) technology represented a major advance in the field of chemogenetics. The original DREADD approach (Roth, 2016) has been superseded by the use of the PSAM-modified ligand-gated ion channels (Sternson and Bleakman, 2020) to activate or silence neurons in response to exogenous agonists, as they allow direct control of ion channels. An early example of the use of ion channels came with the glutamate-gated chloride channels (from *Caenorhabditis elegans*) (Slimko et al., 2002), which can be activated with the anthelmintic drug ivermectin. The immunogenicity of C.elegans –derived ion channels presents a significant obstacle to any clinical use. Compared to these techniques, human ion channels like PSAM^4^ -GlyR are more appealing in terms of developing useful chemogenetic therapies. Therefore, PSAM^4^-glyR was selected for the current study.

PSAM^4^ -GlyR is a chemogenetic receptor with a ligand binding domain formed from a triple mutation in a human α7nAChR and a pore domain from a glycine receptor. This chimeric channel, upon activation by exogenous agonists, shows conductance properties similar to glycine receptors which are chloride-selective. Glycine is one of the primary components that mediate rapid inhibitory neurotransmission. Early experiments showed that glycine diminishes action potential firing by spinal cord neurons. This is thought to occur because once activated, glycine-gated ion channels (GlyRs) lead to hyperpolarization triggered by chloride input or a shunt effect defined by a decrease in membrane resistance following increased chloride conductance (Van den Eynden et al., 2009).

In order to examine chemogenetic neuronal silencing, we selected veratridine as a stimulus to activate DRG neurons. Veratridine, a sodium channel agonist, causes the subsequent activation of voltage-gated calcium channels leading to higher intracellular calcium and increased GCaMP signals (Mohammed et al., 2017). Calcium responses to veratridine have been previously characterized (Mohammed et al., 2017), where veratridine (30 µM) elicited robust responses in roughly 70% of sensory neurons from C57BL/6 mice. We, therefore, selected this concentration for our experiments. We examined the effect of activating PSAM^4^-GlyR with varenicline on the veratridine-elicited calcium responses of DRG neurons using plasmid vector-based transfection (Figure 3), with promising outcomes. Following that, the PSAM^4^-GlyR plasmid was packaged into AAV9, and the system was tested *in vitro*. Results indicated that the application of varenicline (20nM) to PSAM^4^ -GlyR+ DRG neurons completely silenced 80% of them and resulted in approximately 70% reduction in response intensity in the remaining 20% of DRG neurons, without affecting non-transduced cells (Figure 4).

Following successful *in vitro* DRG silencing upon PSAM^4^-GlyR activation, this system was tested *in vivo*. To test the potential of *in vivo* silencing, we relied on GCaMP-based calcium imaging. Through imaging alive, anesthetized transgenic mice carrying calcium indicators (such as GCaMP3), it is possible to observe hundreds to thousands of neurons (Emery et al., 2016; Wang et al., 2018). By examining the patterns of response to mechanical and thermal stimulation of large groups of neurons, this method has provided a useful mechanistic understanding of multiple pain-related conditions such as chemotherapy-induced cold allodynia (MacDonald et al., 2021a; Iseppon et al., 2022). To achieve this, AAV9 viral vectors were used to deliver the PSAM^4^-GlyR transgene to mouse pups that express GCaMP3 under the control of the *Pirt* promoter by an intraplantar injection. The selection of this serotype was based on the fact that AAV9 and AAV2 are the least affected by neutralizing antibodies from mice sera (Rapti et al., 2012). Additionally, previous work from our lab and other labs (Weir et al., 2017) indicates that AAV9 transduces DRG neurons successfully. The intraplantar injection of AAV9 resulted in the successful expression of PSAM^4^-GlyR in the DRG neurons, as demonstrated by the presence of mCherry signal in approximately 70% of the GCaMP3+ DRG neurons (Figure 6 G-I). We also assessed whether the viral administration/receptor expression could alter withdrawal thresholds in the Hargreaves’ test by comparing the viral injected GCaMP3 mice with GCaMP3 mice that were not injected with the virus and no significant difference was detected (figure 6-1B). Following this test, AAV9-PSAM^4^ -GlyR-injected mice were used for *in vivo* calcium imaging. Varenicline treatment (0.3 mg/kg, i.p.) silenced the responses of more than two-thirds of the heat-responding cells in these animals (Figure 5B). Additionally, when mice treated with varenicline were compared with PBS-treated mice, we noticed that varenicline resulted in a significant reduction (∼80%) in the number of heat-responding cells per mm^2^ (Figure 5A). While previous data suggest that the intracellular concentration of chloride ions in the DRG neurons is high and that the opening of these channels could be excitatory (Morales-Aza et al., 2004; Pfeffer et al., 2009; Wood, 2020), our results indicate that the opening of PSAM^4^-GlyR ligand-gated ion channels silenced the responses of DRG neurons to veratridine (30µM) *in vitro* and to heat *in vivo*. These results could be considered unexpected. The mechanisms by which Cl^-^ ion egress may have an inhibitory effect on DRG neurons include two main mechanisms: depolarization block and shunting inhibition (Kaila et al., 2014).

With the successful *in vivo* silencing of DRG neurons upon varenicline-based PSAM^4^-GlyR activation, we decided to test whether silencing can cause an analgesic effect in acute pain behavioral tests, acute inflammatory pain models, and a model of oxaliplatin-induced cold allodynia. For these experiments, AAV9 viral vectors were used to deliver PSAM_4_-GlyR. The results indicated that virally transduced PSAM_4_-GlyR expressing mice exhibit an elevation in their withdrawal thresholds when activated with varenicline, both via intraperitoneal (Figure 6 (A-C)) and intrathecal administration (extended figure 6-1B) and this elevation is reversible after both routes of varenicline administration (extended figure 6-1). For instance, the withdrawal thresholds in the Hargreaves’ test after the intraplantar injection of PGE2 almost tripled in the varenicline-treated mice as opposed to control mice treated with PBS (Figure 6E). Similarly, and unlike the PBS-treated mice, oxaliplatin did not cause cold allodynia in varenicline-treated mice (Figure 6F). Because it was previously reported that oxaliplatin-induced cold allodynia is caused by the unmasking of Na_V_1.8+ silent cold nociceptors, and because the AAV9 virus can transduce all DRG neuronal subsets, we suggest that silent nociceptors expressing Na 1.8 channels were silenced by the PSAM^4^-GlyR (+ varenicline) system (MacDonald et al., 2021a). Because varenicline did not significantly impact the motor abilities of mice that express PSAM^4^-GlyR on a rotarod, the elevation of withdrawal thresholds in behavioral pain tests is unlikely to be caused by an action on motor neurons (Figure 6D). A preprint by Perez-Sanchez et al. showed a similar elevation in withdrawal thresholds in pain assays upon the use of varenicline to silence DRG neurons expressing PSAM^4^-GlyR (Jimena et al., 2023). In their study, Perez-Sanchez et al. also used AAV9 as a vector to induce the expression of PSAM^4^-GlyR in all DRG neurons. They also found impairment of innocuous sensation with viral delivery.

Since viral transduction studies have an intrinsic variability, we designed a mouse line that expresses PSAM^4^-GlyR receptors in a Cre-dependent manner to allow the expression of the PSAM^4^ -GlyR in specific neuronal subsets to enable detailed mechanistic studies. For this mouse line, we relied on the flip-excision (FLEx) switch. For this, our transgene of interest (PSAM^4^-GlyR IRES tdTomato) was inverted and flanked by two antiparallel loxP-type recombination sites. These sites first undergo coding sequence inversion, and then two sites are excised, leaving one of each orthogonal recombination site with an orientation that prevents it from engaging in further recombination. Even though some approaches rely on a single pair of LoxP sites that are arranged in an antiparallel orientation to achieve the inversion of the coding sequence and subsequent cell-specific expression, we decided to adopt this technique as the inversion is unstable and results in a mixture of reverse and forward configurations, and that could lower the expression levels of the transgene of interest (Atasoy et al., 2008). The use of the CAG promoter and the ability to unlock the PSAM^4^-GlyR in any neuronal cell with the use of either the appropriate Cre or the local injection of viruses carrying Cre as a transgene makes this mouse line a valuable research tool that may be applied to any area of research in the neuroscience field.

The established role of Na_V_1.8 in mouse pain has recently been validated by human genetics, as several gain-of-function mutations have been identified in the *SCN10A* gene that encodes for Na_V_1.8 in painful small-fiber neuropathy patients (Faber et al., 2012). Na_V_1.8 channels are primarily expressed in nociceptors (Djouhri et al., 2003; Agarwal et al., 2004; Stirling et al., 2005). The inward flow of sodium ions through Na_V_1.8 channels comprises about (58–90%) of the total influx of sodium during the rising phase of the action potential (Renganathan et al., 2001; Blair and Bean, 2002). Additionally, the conditional ablation of Na_V_1.8+ neurons affects noxious mechanical sensation without impacting innocuous mechanical sensation (Abrahamsen et al., 2008b). Na_V_1.8 thus represents a key target for pain research. We, therefore, selected the Na_V_1.8+ subset of sensory neurons to assess whether an analgesic effect could be seen after silencing these neurons. We explored the role of Na_V_1.8+ neurons in CIBP and neuropathic pain. These models were selected because these conditions are hard to treat. It was found that the bone is one of the most commonly reported cancer pain sites (Barbera et al., 2010; Mayer et al., 2011; Batalini et al., 2017), because of the high prevalence of bone metastases (Coleman, 2001). The prevalence of neuropathic pain in the general population ranges from 3% to 17%. The majority of neuropathic pain medications on the market have a moderate level of effectiveness and have side effects that restrict their use (Cavalli et al., 2019). We, therefore, examined CIBP and neuropathic pain models and the effect of chemogenetic silencing of Na_V_1.8+ neurons. Backed up by previous findings suggesting that Na_V_1.8 channels play an important role in neuropathic pain and CIBP (Roza et al., 2003; Liu et al., 2014), and the analgesic effect of channel knockdown in CIBP (Miao et al., 2010), we tested whether silencing this set can reduce CIBP and pain associated with CCI. Our results indicated that the intraperitoneal administration of varenicline to cancer-bearing mice expressing PSAM^4^-GlyR diminished their weight-bearing deficits, not only highlighting the efficacy of this system but also confirming the involvement of the Na_V_1.8-positive neurons in CIBP (Figure 8A). Similarly, varenicline reversed signs of hypersensitivity (cold, thermal, and mechanical) after sciatic nerve injury in PSAM^4^-GlyR-expressing mice (Figure 9 (A-D)), without affecting innocuous mechanical sensation (figure 7C). These results align with previous findings suggesting the involvement of these neurons in CIBP (Miao et al., 2010) and neuropathic pain (Roza et al., 2003). We also confirmed published data demonstrating that the ablation of Na_V_1.8+ neurons does not impact innocuous touch sensation (Abrahamsen et al., 2008b). The pressing need to find novel analgesics for CIBP and neuropathic pain and the positive outcomes obtained by silencing the Na_V_1.8+ neurons encourage the application of chemogenetic tools for patients living with these conditions if effective viral targeting can be established. In addition, these findings highlight the role of this neuronal subset in both neuropathic pain and CIBP, helping to identify targets within this subset for future clinical application.

Our findings thus show that the activation of PSAM_4_-GlyR in DRG neurons silences them *in vitro* and *in vivo*, as demonstrated by the profound reduction in calcium signals within the DRG neurons after stimulation. When this system is tested in live and behaving animals, and after the expression of PSAM^4^-GlyR in all DRG neuronal subsets by viral vectors, an elevation of withdrawal thresholds in various acute pain behavioral tests was seen. Additionally, upon varenicline administration, these mice showed a significant reduction in inflammatory-driven heat hyperalgesia as well as oxaliplatin-induced cold allodynia. While these effects are beneficial, the lack of specificity in PSAM^4^-GlyR expression resulted in significant impairment of innocuous mechanical sensation. On the other hand, when a mouse line was used to restrict the expression of PSAM_4_-GlyR to Na_V_1.8+ neurons, a reduction in acute noxious mechanical and thermal sensation was detected without a significant effect on innocuous mechanical sensation. This mouse line was then tested in various chronic pain models, such as cancer-induced bone pain and neuropathic pain, and a reversal of signs of mechanical, thermal, and cold allodynia was seen. The reversible neuronal silencing that can be achieved with chemogenetic tools makes them an attractive way of targeting neuronal subsets, which could help in complex conditions like cancer-induced bone pain and neuropathic pain that suffer from high attrition rates in the clinic. The fact that we have shown that with viral delivery, PSAM^4^ -GlyR was expressed in the DRG neurons and could be activated exogenously to lead to favorable and reversible behavioral outcomes, makes PSAM^4^ -GlyR a potential therapeutic tool. Methods to restrict the expression of the receptor to certain neuronal subsets, such as the Na_V_1.8+ neurons are desirable but not presently available. Cre-dependent reversible neuronal silencing with the FLEx PS AM^4^ - GlyR mouse is nonetheless an effective research tool, applicable to a range of neurobiological questions.

## Acknowledgments

We thank Dr Federico Iseppon, Dr Naxi Tian and Alexander Fudge for their valuable contributions. The graphical abstract, Figure 1, and Figure 2 were generated using BioRender.com. pCAG PSAM4 GlyR IRES EGFP was a gift fr m Scott Sternson (Addgene plasmid # 119739 ; http://n2t.net/addgene:119739 ; RRID:Addgene_119739).

## Extended Data

**Figure 6-1:**
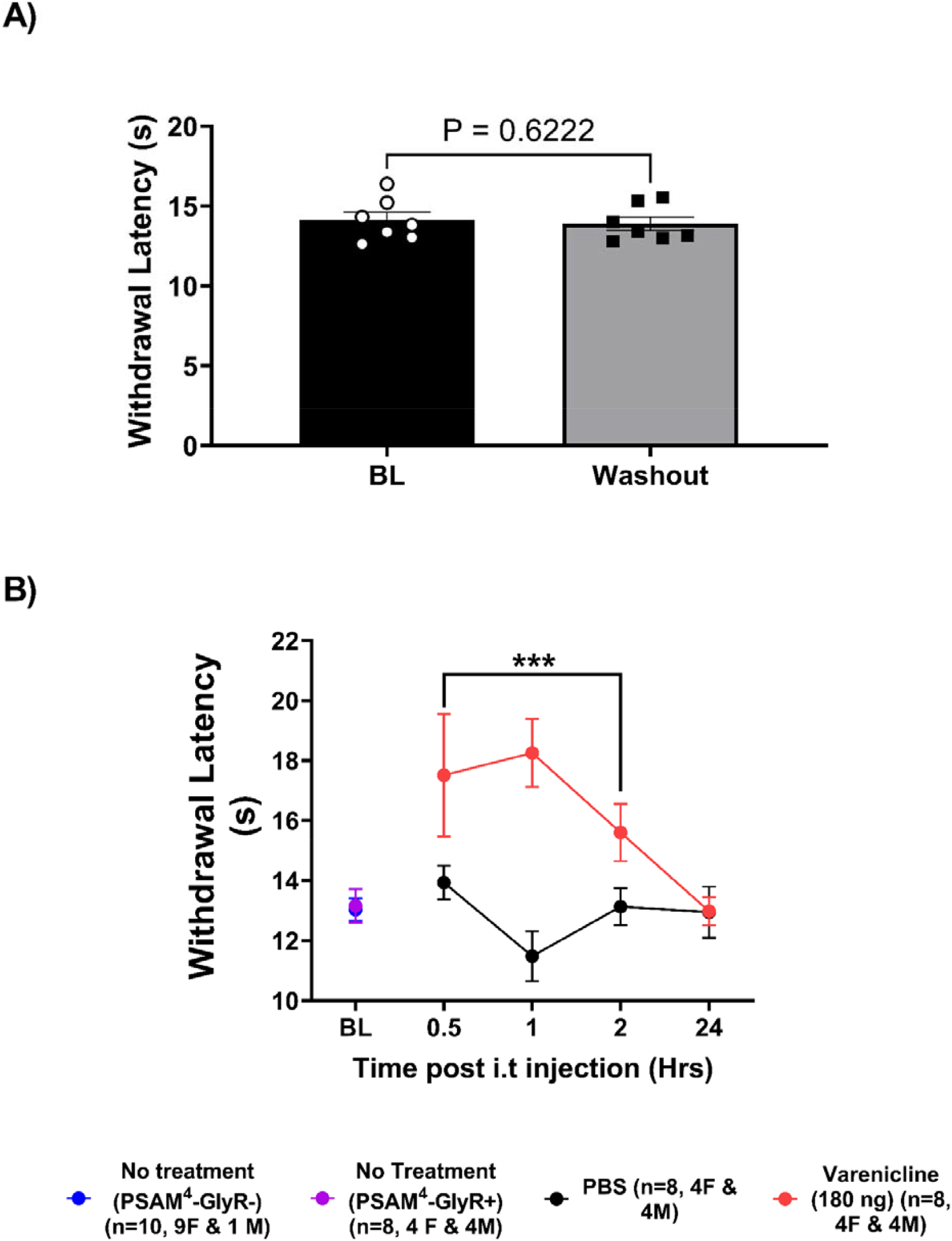
Varenicline causes a reversible increase in the withdrawal threshold of mice expressing PSAM^4^ -GlyR in the Hargreaves’ test. A. Mice expressing PSAM^4^ -GlyR and treated with varenicline (0.3 mg/kg, i.p) experience an elevation in their withdrawal threshold in the Hargreaves’ test and then return to their baseline thresholds 6 hours post-treatment (washout). N=7 (4M & 3F). B. While the expression of PSAM^4^ -GlyR using viral vectors in GCaMP3 mice did not cause a significant change in the withdrawal thresholds in the Hargreaves’ test compared to GCaMP3 mice without PSAM^4^ -GlyR (p-value =0.8485, unpaired t-test, Blue vs. purple), the intrathecal administration of varenicline to PSAM^4^ -GlyR expressing GCaMP3 mice elevated their withdrawal thresholds significantly, as indicated by statistical analyses using the two-way ANOVA test to compare varenicline and PBS-treated PSAM^4^ -GlyR-expressing mice in the period between 0.5- and 2 hours post-treatment (p-value= 0.0006, red vs. black). Mice returned to their baseline withdrawal threshold 24 hours post-treatment with intrathecal varenicline (the paired t-test between baseline without treatment and 24 hours post-varenicline, p-value= 0.6846). Error bars represent SEM. PBS: Phosphate-buffered saline, BL: baseline.

